# Variation in supplemental carbon dioxide requirements defines lineage-specific antibiotic resistance acquisition in *Neisseria gonorrhoeae*

**DOI:** 10.1101/2022.02.24.481660

**Authors:** Daniel H.F. Rubin, Kevin C. Ma, Kathleen A. Westervelt, Karthik Hullahalli, Matthew K. Waldor, Yonatan H. Grad

## Abstract

The evolution of the obligate human pathogen *Neisseria gonorrhoeae* has been shaped by selective pressures from diverse host niche environments^1,2^ as well as antibiotics^3,4^. The varying prevalence of antibiotic resistance across *N. gonorrhoeae* lineages^5^ suggests that underlying metabolic differences may influence the likelihood of acquisition of specific resistance mutations^6,7^. We hypothesized that the requirement for supplemental CO_2_, present in approximately half of isolates^8^, reflects one such example of metabolic variation. Here, using a genome-wide association study and experimental investigations, we show that CO_2_-dependence is attributable to a single substitution in a β-carbonic anhydrase, *canB*. CanB^19E^ is necessary and sufficient for growth in the absence of CO_2_, and the hypomorphic CanB^19G^ variant confers CO_2_-dependence. Furthermore, ciprofloxacin resistance is correlated with CanB^19G^ in clinical isolates, and the presence of CanB^19G^ increases the likelihood of acquisition of ciprofloxacin resistance. Together, our results suggest that metabolic variation has impacted the acquisition of fluoroquinolone resistance.

## Main Text

*N. gonorrhoeae* is commonly cultured in the presence of supplemental CO_2_^9^, though the molecular basis for this trait has not been elucidated. We first investigated the mechanism of CO_2_-dependence using a genome-wide association study (GWAS). To generate phenotypes for GWAS, clinical and laboratory strains of *N. gonorrhoeae* were assessed for growth in the presence and absence of supplemental CO_2_. The GWAS revealed a significant association with a single genomic site, a missense mutation in *ngo2079* (**Fig. 1A**). In a parallel unbiased method, genomic DNA from FA6140, an *N. gonorrhoeae* strain that does not depend on supplemental CO_2_, was used to transform FA19, a lab isolate that requires supplemental CO_2_ (**Supp. Fig. 1A**). Transformants that grew in the absence of supplemental CO_2_ had acquired multiple SNPs in the region of *ngo2079* (**Supp. Fig. 1B, C**), including the SNP identified by GWAS.

**Figure 1.**
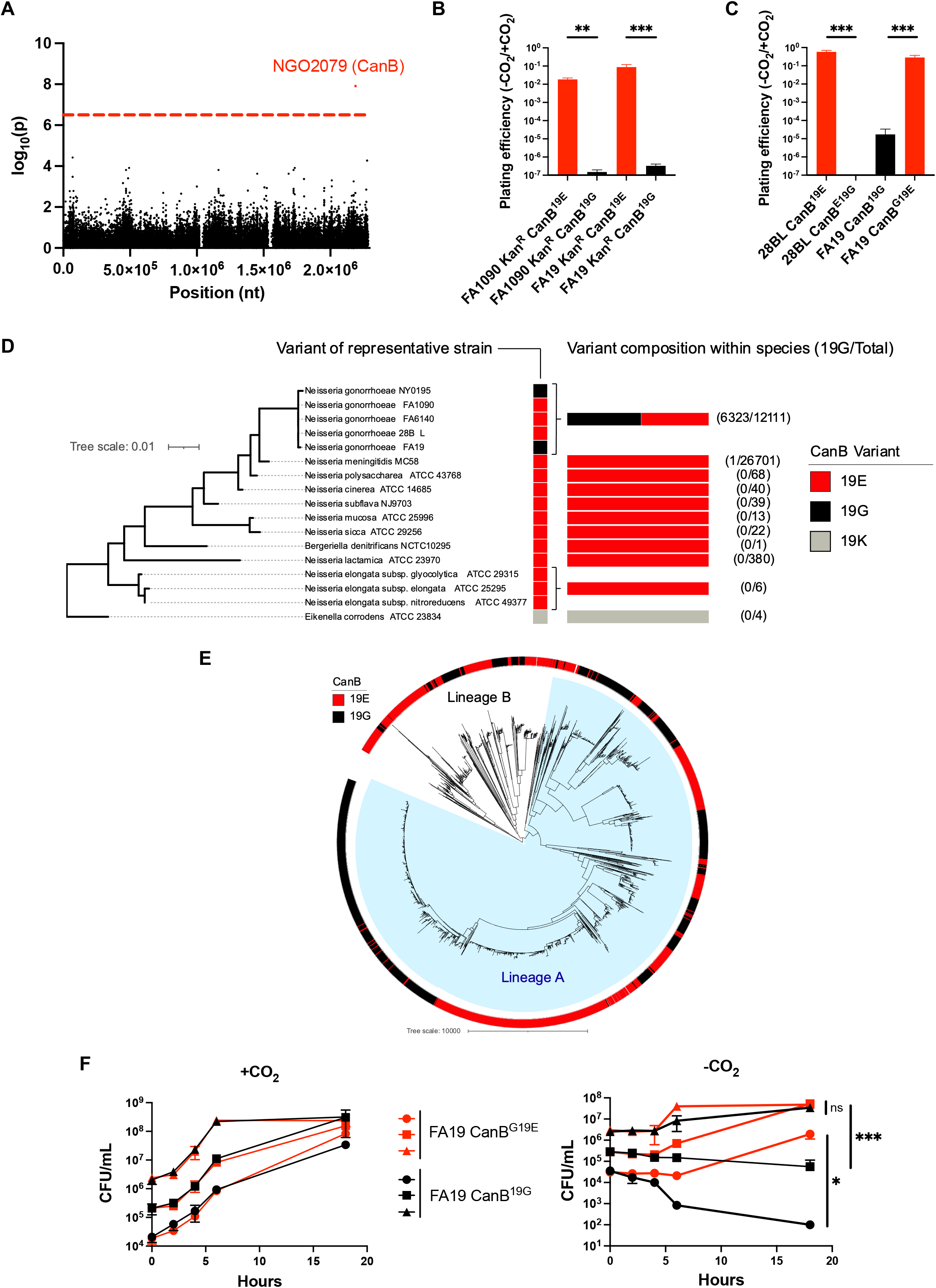
A SNP in *ngo2079*, encoding a β-carbonic anhydrase, is necessary and sufficient to explain dependence on supplemental CO_2_ in *N. gonorrhoeae*. **(A)** Manhattan plot of SNPs associated with dependence on supplemental CO_2_ across 30 strains of *N. gonorrhoeae*. Dotted line represents a p-value of 0.05 with Bonferroni correction for 57,691 unitigs. **(B)** Plating efficiencies in the absence and presence of 5% supplemental CO_2_ of CanB variants introduced by kanamycin co-selection in *N. gonorrhoeae* strains FA1090 (parental CanB^19E^) and FA19 (parental CanB^19G^) (N ≥ 6 from two independent experiments). Significance determined by Mann-Whitney U test. **(C)** Plating efficiencies as in (**B**) of *N. gonorrhoeae* strains 28BL (parental CanB^19E^) and FA19 (parental CanB^19G^) with isogenic CanB mutants (N ≥ 6 from two independent experiments). Significance determined by Mann-Whitney U test. **(D)** NGO2079/CanB variants across sequenced *Neisseria* and related species represented alongside a 16S maximum-likelihood tree. Branch length represents number of substitutions per site. **(E)** Maximum-likelihood phylogenetic tree of 5,007 strains of *N. gonorrhoeae* with a colored track representing CanB variant. Branch length represents total number of substitutions after removal of predicted recombinations. Lineage A represents isolates that arose phylogeographically After the introduction of *N. gonorrhoeae* into Asia, while Lineage B represents isolates that arose Before that breakpoint, as defined by Sánchez-Busó *et al*.^5^ **(F)**Growth of FA19 CanB isogenic strains in the presence and absence of supplemental CO_2_ (N=3, representative of two independent experiments). Significance at 18 hours determined by unpaired two-sided t-test. *p<0.05, **p<0.01, ***p<0.001

The *ngo2079* c.56G→A SNP encodes a G→E mutation at amino acid 19 of NGO2079, a β-carbonic anhydrase. *N. gonorrhoeae* contains two additional carbonic anhydrases: a well-characterized essential periplasmic ɑ-carbonic anhydrase^10-12^ and a putative nonessential cytosolic ɣ-carbonic anhydrase^13^. All carbonic anhydrases catalyze the conversion of H_2_O and gaseous CO_2_ to aqueous bicarbonate. Aqueous bicarbonate is then used as a carbon building block in multiple pathways within the cell, including the synthesis of TCA cycle components, nucleotides, and phospholipids^14^ (**Supp. Fig. 1C**). As *ngo2079* is the sole predicted *N. gonorrhoeae* β-carbonic anhydrase, we named this gene *canB* to parallel the corresponding carbonic anhydrase *can* in *E. coli*. The SNP is located outside of the highly conserved carbonic anhydrase domain (**Supp. Fig. 1D**) and does not appear to be within a signal sequence domain by SignalP^15^. Structures from Alphafold^16^ indicate that the variant amino acid lies between two N-terminal alpha-helices, but it does not appear at the dimerization interface or the characterized active site (**Supp. Fig. 1E**).

We validated our GWAS results by introducing the CanB variants into *N. gonorrhoeae*. Replacement by kanamycin co-selection of the CanB^19E^ in FA1090 with CanB^19G^ and of the CanB^19G^ in FA19 with CanB^19E^ showed that dependence on supplemental CO_2_ was attributable to CanB^19G^ (**Fig. 1B**), while similar results for single point mutants in 28BL (19E→G) and FA19 (19G→E) (**Fig. 1C**) support this idea. These results indicate that CanB^19E^ is both necessary and sufficient for growth in the absence of supplemental CO_2_. The concentration of CO_2_ that differentiates this dependency is between atmospheric (∼410 ppm) and 0.1% CO_2_ (1000 ppm) (**Supp. Fig. 2A**), slightly lower than has been found in other bacteria^17^ and considerably below the 5% CO_2_ standardly used in *N. gonorrhoeae* growth protocols^9^.

CanB^19G^, the allele that confers CO_2_-dependence, appears to have arisen multiple times in *N. gonorrhoeae*, whereas genomes from *Neisseria spp*. and other closely related bacteria encode only CanB^19E^ (**Fig. 1D**). Analysis of the PubMLST database of more than 36,000 *Neisseria spp*. genomes showed that only one non-gonococcal *Neisseria* isolate carries CanB^19G^, a urethral isolate of *N. meningitidis* (PubMLST id: 82427) that appears to have a acquired the *canB* region from *N. gonorrhoeae* (100% identity to sequenced *N. gonorrhoeae* isolates). Within *N. gonorrhoeae*, ∼50% of sequenced isolates from across the phylogenetic tree have CanB^19G^ (**Fig. 1D, E**). The proportion of sequenced isolates with CanB^19G^ has remained relatively stable at ∼50% during last twenty years (**Supp. Fig. 6A**), though this allele appears to have been less common in the pre-antibiotic era of the 1930s (9% of pre-1930 *N. gonorrhoeae* isolates from the PubMLST database).

Although CanB^19E^ provides a growth advantage in the absence of supplemental CO_2_, the growth of FA19 CanB^19G^ and CanB^G19E^ in the presence of supplemental CO_2_ was indistinguishable and independent of inoculum size (**Fig. 1F, left**). Furthermore, in the presence of CO_2_, kanamycin-labeled CanB isogenic FA19 strains remained at equal proportions in competition (**Supp. Fig. 2B**). The CanB^19G^ variant, therefore, does not confer a general fitness defect but a specific growth disadvantage in the absence of CO_2_. As shown in **Fig. 1F, right**, this disadvantage is concentration-dependent and is exacerbated at lower starting inocula. Although FA19 CanB^19E^ is able to grow at all starting CO_2_ concentrations, both strains have a substantially longer lag phase in the absence of CO_2,_ suggesting that intracellular bicarbonate plays a critical role in lag phase, similar to *N. meningitidis*^*18*^. This competitive disadvantage is not rescued by extracellular sodium bicar-bonate or oxaloacetate, a downstream product of intracellular bicarbonate (**Supp. Fig. 2C**). Thus, in contrast to many bacteria that are able to acquire bicarbonate from the environment via inorganic carbon transporters^19^, *N. gonorrhoeae* does not appear to have a CO_2_-concentrating mechanism apart from cytosolic carbonic anhydrases, explaining why exogenous bicarbonate does not rescue the competition defect. Supplementation with metabolites downstream of intracellular bicarbonate (**Supp. Fig. 1D**), such as pyruvate, sub-minimum inhibitory concentration (MIC) palmitic acid, and purines were also unable to rescue this defect (**Supp. Fig. 2D**).

*E. coli* MG1655 was used to compare the activity of the CanB variants. Deletion of one of the two endogenous *E. coli* β-carbonic anhydrases, *can*, rendered MG1655 capnophilic and dependent on supplemental CO_2_ (**Fig. 2A**, top two quadrants). Complementation with IPTG-inducible *N. gonorrhoeae* CanB variants restored growth in the absence of supplemental CO_2_. However, lower concentrations of IPTG were required for complementation with CanB^19E^ versus CanB^19G^ (**Fig. 2B**, lower two quadrants), even though the abundance of CanB^19E^ and CanB^19G^ were similar at equivalent levels of induction (**Supp. Fig. 3A**), suggesting that CanB^19G^ retains some enzymatic activity but does not restore growth as efficiently as CanB^19E^.

**Figure 2.**
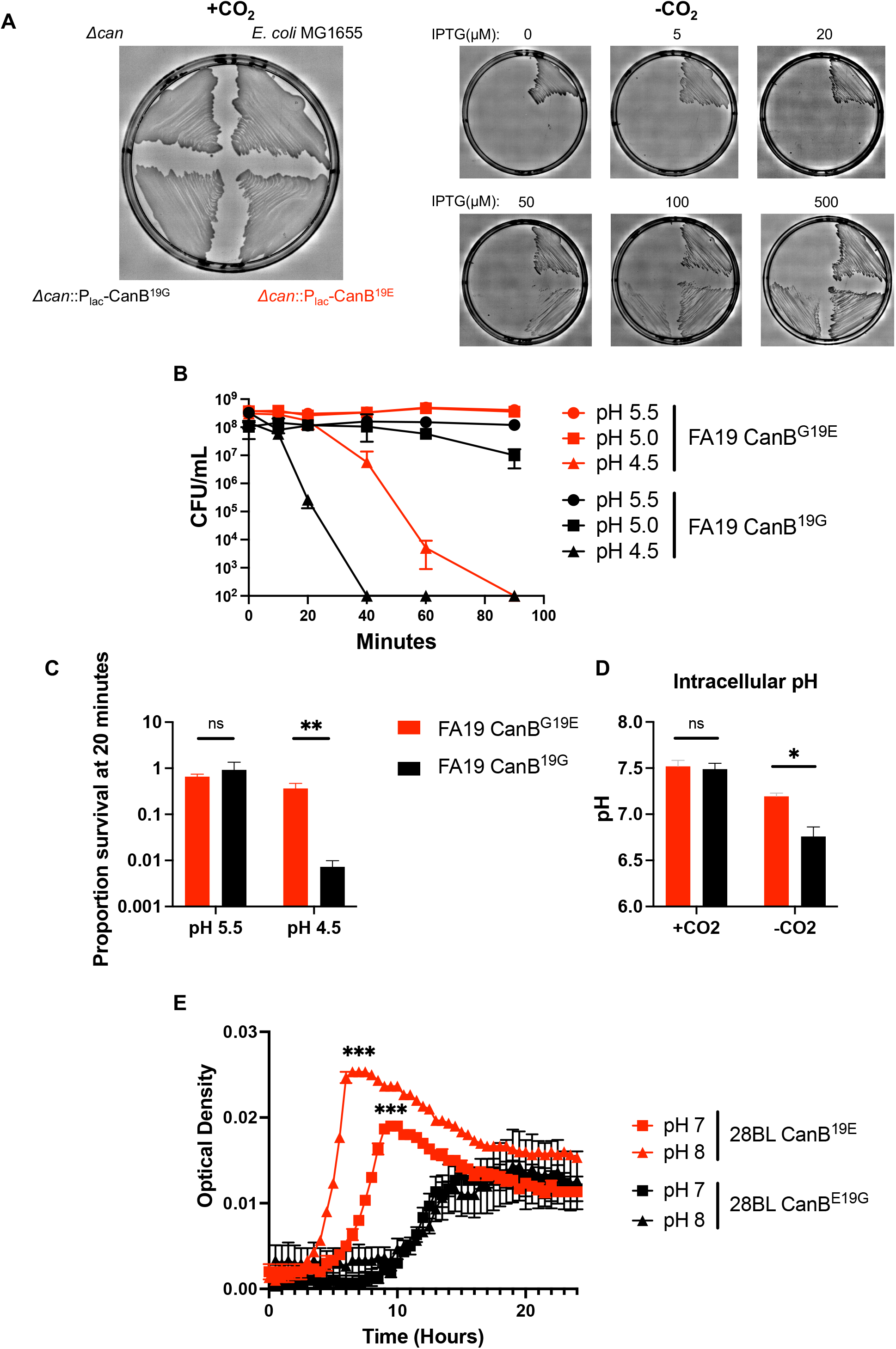
The 19G variant of CanB is a functional hypomorph. **(A)** Growth of parental *E. coli* MG1655, MG1655Δ*can*, and MG1655Δ*can* complemented with CanB variants under different levels of induction. **(B)** Survival curve under multiple low pH conditions of FA19 isogenic CanB isolates in monoculture (N=3, representative of three independent experiments). **(C)** As in (**B)**, survival at 20 minutes of exposure to low pH conditions (N=6, from two independent experiments). Significance by Mann-Whitney U test. **(D)** Intracellular pH of FA19 CanB^19G^ and FA19 CanB^G19E^ as measured by flourimetric dye in the presence and absence of supplemental CO_2_ (N=3, representative of two independent experiments). **(E)** Growth of the 28BL CanB isogenic pair in liquid culture under anaerobic conditions (N=3, representative of two independent experiments). Significance by unpaired two-sided t-test. *p<0.05, **p<0.01, ***p<0.001

Since CanB^19G^ is a hypomorph, we hypothesized that the CanB variants result in altered gonococcal physiology. At pH 4.5, an acidic condition similar to healthy vaginal pH^20^, FA19 CanB^19G^ died more quickly than the isogenic FA19 CanB^19E^ (**Fig. 2B, C**), consistent with the finding that, in the absence of supplemental CO_2_, FA19 CanB^19G^ is less able to buffer intracellular pH (**Fig. 2D**). The CanB^19E^ survival advantage at low pH did not extend, however, to intracellular survival within immortalized macrophages (**Supp. Fig. 4A**). At higher pH and in aerobic conditions, neither the CanB^19E^ or CanB^19G^ variant displayed a substantial advantage (**Supp. Fig. 2B, Supp. Fig. 4B**). In contrast, in anaerobic conditions, FA19 CanB^19E^ had a distinct growth advantage across the pH range supportive of growth (**Fig. 2E, Supp. Fig. 4C**). Our data thus show that the CanB^19E^ allele provides an advantage at low pH and under anaerobic conditions, suggesting that CanB^19E^ may allow for buffering in the acidic vaginal environment.

Carbonic anhydrases have been previously shown to be targets of sulfa antibiotics^12^ and to mediate natural competence in *N. meningitidis*^21^. Sulfa antibiotics were one of the first antibiotics to be used to treat gonorrhea^22^ and primarily target dihydropteroate synthase in the folate synthesis pathway. Both CanB variants had similar sulfamethoxazole and trimethoprim/sulfamethoxazole MICs across two genetic backgrounds (**Supp. Table 2A**). Furthermore, MG1655Δ*can* complemented with CanB^19G^ or CanB^19E^ also had similar sulfamethoxazole MICs (**Supp. Table 2B**). The CanB variants therefore do not have different susceptibility to sulfa antibiotics. With respect to competence, a *canB* knockout was dependent on supplemental CO_2_ (**Supp. Fig. 5A**), but both the wild-type and knockout had the same level of natural competence, as measured by the introduction of a single SNP in *gyrB* conferring nalidixic acid resistance (**Supp. Fig. 5B**). Similarly, competence was not associated with the CanB isogenic pairs of FA19 and 28BL (**Supp. Fig. 5B**). Thus, selection for CanB variants does not appear to be driven by associations with sulfa susceptibility or natural competence.

In *N. gonorrhoeae*, antibiotic resistance varies across the phylogenetic tree, with some lineages displaying more resistance than others, though the reasons explaining these differences are not known. One hypothesis is that lineages may be subject to demographically or geographically variable antibiotic pressures. The overrepresentation of heterosexuals in Lineage B and men-who-have-sex-with-men in Lineage A lend support to this hypothesis^5^ (**F**i**gure 1E**), but the rapid global spread of *N. gonorrhoeae* and the extensive bridging across sexual networks argue against it^23^. Alternatively, we hypothesized that genomic and associated metabolic variation could modify the likelihoods of acquiring and maintaining resistance. Support for this idea comes from previous research showing that intracellular metabolite perturbations can shape the likelihood of developing resistance^24,25^. To begin to address whether the CanB variants have shifted the evolutionary landscape of *N. gonorrhoeae* antibiotic resistance, we analyzed whether the variants are associated with antibiotic resistance on the population level. We found that the CanB^19G^ variant is associated with >4-fold higher MICs for ciprofloxacin (average log_2_ MIC 0.43 vs. 0.10 µg/mL) (**Fig. 3A**), whereas MICS for azithromycin (0.32 µg/mL vs. 0.24 µg/mL), penicillin (0.41 vs. 0.70 µg/mL), and ceftriaxone (.0093 µg/mL vs. .013 µg/mL) did not differ (**Supp. Fig. 6B**). CanB^19G^ is also associated with the most common resistance determinant for ciprofloxacin, GyrA^91F^, as 52.3% of CanB^19G^ isolates have GyrA^91F^, as compared to only 30.4% of CanB^19E^ isolates (p<0.0001, Pearson’s chi-square) (**Fig. 3B**). This relationship holds true across years, as the proportion of GyrA^91F^ and CanB^19G^ isolates per year in the United States have tracked closely since 2000 (**Fig. 3C**). On a country-by-country level, isolates with CanB^19G^ have a comparatively higher ciprofloxacin MIC (**Supp. Fig. 6C**), indicating that the association is true across datasets and geography. The simplest explanation is that CanB^19G^ directly increases ciprofloxacin resistance. However, the isogenic CanB FA19 strains have comparable levels of resistance to ciprofloxacin (**Supp. Table 2C**).

**Figure 3.**
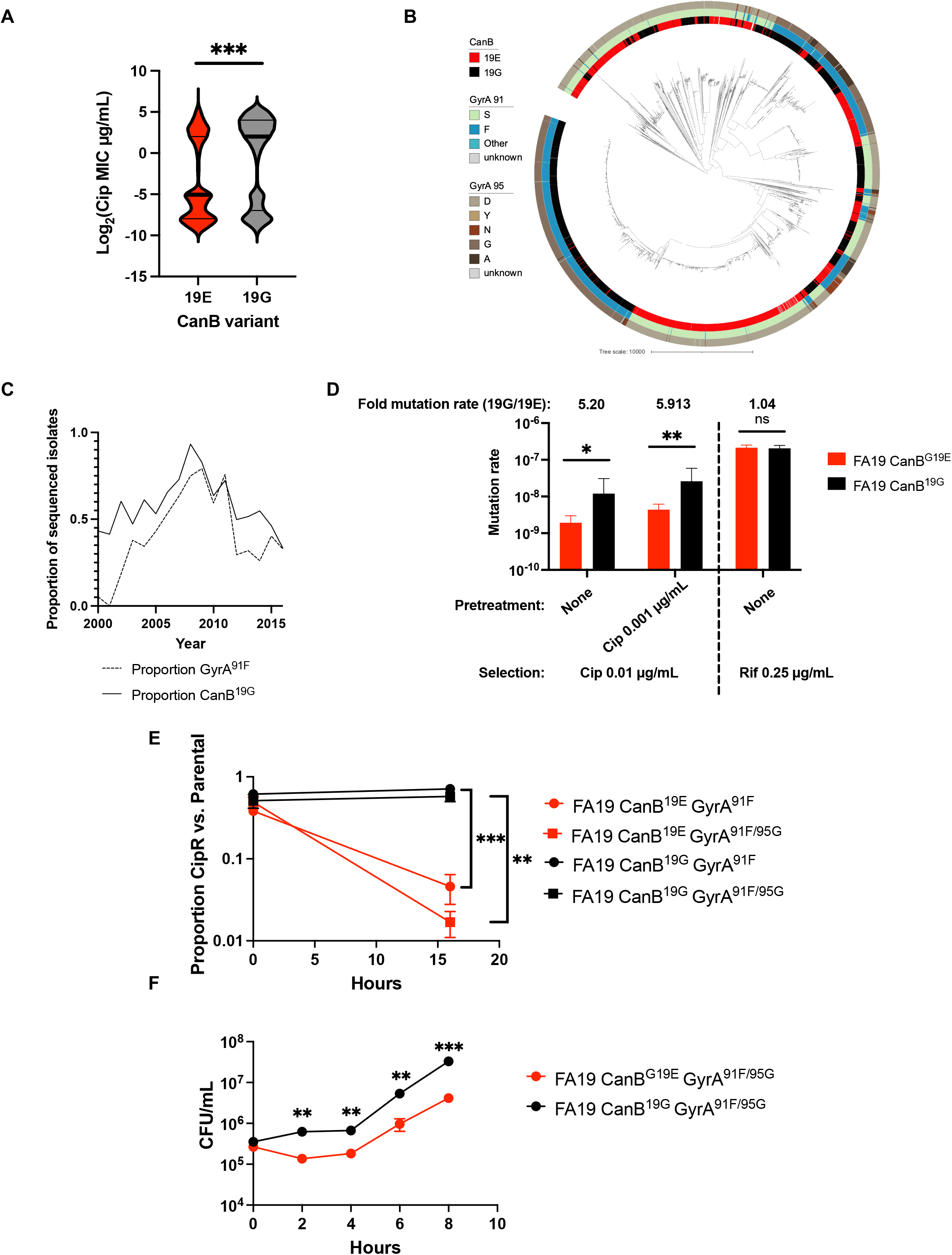
The CanB^19G^ variant facilitates acquisition of ciprofloxacin resistance. **(A)** Violin plot of ciprofloxacin MICs for 8,912 *N. gonorrhoeae* clinical isolates. Thick lines represent median, thin lines represent 25^th^/75^th^ percentiles. Significance by Mann-Whitney U test. **(B)** Maximum-likelihood tree, as in **Figure 1C**, overlaid with tracks representing ciprofloxacin conferring variants of GyrA. **(C)** Proportion of sequenced U.S. isolates stratified by year that have the CanB^19G^ variant and the ciprofloxacin resistance-conferring GyrA^91F^ variant. **(D)** Calculated mutation rates to ciprofloxacin and rifampin by fluctuation analysis of isogenic CanB strains +/- pretreatment with sub-MIC ciprofloxacin. Significance by unpaired two sample t-test (N=144, representative of two independent experiments). **(E)** Competition between ciprofloxacin resistant isogenic CanB strains and susceptible parental strains (N=3, representative of two independent experiments). Significance by unpaired two sample t-test. **(F)** Growth curves of CanB isogenic FA19 strains with ciprofloxacin resistance-determining *gyrA* alleles (N=3, representative of two independent experiments). Significance by unpaired two sample t-test. *p<0.05, **p<0.01, ***p<0.001

We therefore theorized that the CanB^19G^ allele may lead to a greater rate of acquisition of ciprofloxacin-specific resistance. A fluctuation assay to estimate the mutation rate to different antibiotics was used to test this idea. FA19 CanB^19G^ had a significantly higher mutation rate than FA19 CanB^19E^ when plated on ciprofloxacin (**Fig. 3D**). Furthermore, the fold-change in mutation rate remained similar following preincubation with sub-MIC ciprofloxacin, which is known to increase the rate of mutation and acquisition of resistance^26^. This suggests that it is not exposure to ciprofloxacin itself that mediates the difference between CanB^19G^ and CanB^19E^, but rather a difference in the pre-exposure populations. There was no difference in killing by ciprofloxacin for either strain, providing further evidence that the difference in mutation rate is not due to differential viability in the presence of drug (**Supp. Fig. 7A**).

We reasoned that there were two potential explanations for the differences in ciprofloxacin mutation rates associated with the CanB^19G^ and CanB^19E^ variant strains. The first is that there is a difference in basal mutation rate between the strains, manifesting as a higher rate of resistance to all antibiotics. This does not appear to be the case, since isogenic CanB FA19 strains had very similar mutation rates to rifampin, an antibiotic that targets RNA polymerase (**Fig. 3D**). Furthermore, a similar fluctuation assay in *E. coli* showed that induction of CanB^19G^ led to a higher rate of ciprofloxacin-specific resistance acquisition (**Supp. Fig. 3B**).

A second possibility is that the CanB^19G^ strains have an increased ability to tolerate *gyrA* mutations. If this were true, we would expect that CanB^19E^ *gyrA* mutants would have a relative competitive disadvantage compared to the parental strains. To investigate this possibility, we generated GyrA^91F^ and GyrA^91F/95G^ mutants in both FA19 CanB backgrounds. For both the GyrA^91F^ and GyrA^91F/95G^ mutants, the CanB^19E^ strain was significantly disadvantaged against the parental strain (**Fig. 3E**). In monoculture, FA19 CanB^19E^ GyrA^91F/95G^ also grew significantly slower than FA19 CanB^19G^ GyrA^91F/95G^ (**Fig. 3F**), as did the equivalent mutants in the 28BL background (**Supp. Fig. 7B**). We generated spontaneous ciprofloxacin resistant mutants, all of which bore GyrA^95^ variants seen in clinical isolates (**Supp. Table 2D**). The ciprofloxacin-resistant FA19 CanB^19G^ strains competed better than ciprofloxacin-resistant CanB^19E^ strains against the susceptible parental strains (**Supp. Fig. 7C**) and showed faster growth in monoculture (**Supp. Fig. 7D**). Taken together, these results suggest that the increased rate of acquisition of ciprofloxacin resistance in the CanB^19G^ background is due to its higher relative fitness in the background of gyrase mutations.

Collectively, our findings suggest that CanB represents a novel metabolic mediator of drug resistance acquisition. Although *canB* is not essential for *in vitro* growth (**Supp. Fig. 5A**), we observed no evidence of premature stop codons in a collection of >16,000 genomes from clinical isolates, indicating its important role *in vivo*. The hypomorphic CanB^19G^ allele is present almost exclusively in *N. gonorrhoeae* among the *Neisseriae* and can be found in samples predating the introduction of antibiotics, suggesting it reflects adaptation to the urogenital niche or sexual transmission. CanB^19E^ also shows lineage-specific background dependence, as the alleviation of CO_2_-dependence in clinical isolate NY0195 requires further introduction of AasN^Q321E^, a mutation in an acyl-ACP synthetase involved in fatty acid uptake^27^ (**Supp. Fig. 8A**). The AasN^321Q^ variant is restricted to a single lineage that carries the ceftriaxone-resistance determining PenA34 allele (**Supp. Fig. 8B**), suggesting further complex connections between central metabolism and antibiotic resistance.

CanB^19G^ is therefore an example of a widespread variant that facilitates acquisition of drug-specific resistance through shifts in the gonococcal metabolic landscape and without directly affecting drug susceptibility or tolerance. This finding expands upon recent work showing that mutations in important metabolic genes can change the drug-specific resistance profile of clinical isolates.^6^ By showing how specific lineages are more likely to acquire specific resistance determinants, our work provides a new perspective for understanding the evolution of antibiotic resistance.

## Author contributions

DHFR and KCM performed the GWAS and statistical analyses. DHFR and KAW performed the anaerobic experiments. DHFR and KH performed the macrophage experiments. The remainder of the experimental work was performed by DHFR. All authors (DHFR, KCM, KAW, KH, MKW, YHG) contributed to data interpretation. YHG supervised and managed the study. DHFR and YHG wrote the manuscript. All authors reviewed and edited the final manuscript. All authors were responsible for the decision to submit for publication.

## Acknowledgements

We thank the members of the Grad Lab and the Waldor lab for their invaluable feedback into this project; Samantha Palace in particular for her thoughts on experimental design; and the Microbial Genome Sequencing Center (https://www.migscenter.com/) for their work sequencing strains. This work was supported by NIH R01 AI132606 and R01 AI153521 and by the Smith Family Foundation Odyssey award (YHG) and R01 AI 042347-24 (MKW). DHFR is funded by NIH F30 AI160911-01 and NIH T32 GM007753. KH is supported by NIH F31 AI156949-01.

