## Supplementary Material for "Variation in supplemental carbon dioxide requirements defines lineage-specific antibiotic resistance acquisition in *Neisseria gonorrhoeae*"

Supplementary Material for Rubin *et al.*
Supplementary Figures 1-8
Supplementary Tables 1 and 2
Methods and Materials

| Plasmid/Strain | Property(ies) | Citation |
| --- | --- | --- |
| **Plasmids** | | |
| pDR1 | $Kan^R$ derivative of pKH37 | This study |
| pDR35 | pUC19 with $Kan^R$ cassette to knockout *canB* | This study |
| pDR42 | pMR33 with $CanB^{19G}$-FLAG, cloned into SacI/XbaI site | This study |
| pDR43 | pMR33 with $CanB^{19E}$-FLAG, cloned into SacI/XbaI site | This study |
| pDR46 | pUC19 with $Cm^R$ cassette to knockout *aasN* | This study |
| pDR48 | pDR75 with $AasN^{321Q}$, cloned into PacI/XbaI site | This study |
| pDR49 | pDR75 with $AasN^{321E}$, cloned into PacI/XbaI site | This study |
| pDR50 | pUC19 wih $Kan^R$ cassette to co-select for $CanB^{19E}$ | This study |
| pDR51 | pUC19 wih $Kan^R$ cassette to co-select for $CanB^{19G}$ | This study |
| pDR55 | pRE107 with homology to knockout *can* in MG1655 | This study |
| pDR75 | $Kan^R$ derivative of pMR32 | This study |
| pKH37 | $Cm^R$, lac-inducible, inserts into *lctP*/*aspC* site of *N. gonorrhoeae* | [1] |
| pMR32 | $Cm^R$, constitutive *opa* promoter, inserts into *trpB*/*iga* site of *N. gonnorrhoeae* | [2] |
| pMR33 | $Cm^{R,}$ lac-inducible, inserts into *trpB*/*igA* site of *N. gonnorrhoeae* | [2] |
| pUC19 | $Amp^R$, for cloning in *E. coli* | [3] |
| pRE107 | $Amp^R$, for allelic exchange in *E. coli* | [4] |
| **Strains** | | |
| FA19 | 1962 *N. gonorrhoeae* isolate from disseminated infection, widely used laboratory strain | [5] |
| FA1090 | Pre-1981 *N. gonorrhoeae* isolate, widely used laboratory strain | [6] |
| FA6140 | 1983 *N. gonorrhoeae* isolate, widely used laboratory strain | [7] |
| 28BL | 1974 *N. gonorrhoeae* isolate from disseminated infection, widely used laboratory strain | [8] |
| MG1655 | *E. coli* laboratory strain derived from K-12 | [9] |
| GCGS0457 | *N. gonorrhoeae* clinical isolate | [10] |
| FA19_FA6140gDNA_1 | FA19 transformed with gDNA from strain FA6140 and selected in the absence of supplemental $CO_2$, isolate 1 | This study |

| | | |
|---|---|---|
| FA19_FA6140gDNA_2 | FA19 transformed with gDNA from strain FA6140 and selected in the absence of supplemental $CO_2$, isolate 2 | This study |
| FA19 pDR50 | FA19, co-selected for $CanB^{19E}$ with $Kan^R$ cassette | This study |
| FA19 pDR51 | FA19, co-selected for $CanB^{19G}$ with $Kan^R$ cassette | This study |
| FA1090 pDR50 | FA1090, co-selected for $CanB^{19E}$ with $Kan^R$ cassette | This study |
| FA1090 pDR51 | FA1090, co-selected for $CanB^{19G}$ with $Kan^R$ cassette | This study |
| 28BL $CanB^{E19G}$ | 28BL selected for dependence on supplemental $CO_2$ with targeted insertion of $CanB^{19G}$ | This study |
| FA19 $CanB^{G19E}$ | FA19 selected for growth in the absence of supplemental $CO_2$ with targeted insertion of $CanB^{19E}$ | This study |
| FA19 pDR1 | FA19 marked with $Kan^R$ via pDR1 | This study |
| FA19 $CanB^{G19E}$ pDR1 | FA19 $CanB^{G19E}$ marked with $Kan^R$ via pDR1 | This study |
| MG1655Δ*can* | MG1655 with deletion of *can* by allelic exchange selection with pDR55 | This study |
| MG1655Δ*can* pDR42 | MG1655Δ*can* with lactose-inducible expression of $CanB^{19G}$-FLAG | This study |
| MG1655Δ*can* pDR42 | MG1655Δ*can* with lactose-inducible expression of $CanB^{19E}$-FLAG | This study |
| NY0195 $CanB^{G19E}$ | NY0195 selected for growth in the absence of supplemental $CO_2$ with targeted insertion of $CanB^{19E}$ | This study |
| NY0195 $CanB^{E19G}$_GCGS0457gDNA_1 | NY0195 $CanB^{E19G}$ transformed with gDNA from strain GCGS0457 and selected in the absence of supplemental $CO_2$, isolate 1 | This study |
| NY0195 $CanB^{E19G}$_GCGS0457gDNA_2 | NY0195 $CanB^{E19G}$ transformed with gDNA from strain GCGS0457 and selected in the absence of supplemental $CO_2$, isolate 2 | This study |
| NY0195 $CanB^{G19E}$ $AasN^{Q321E}$ | NY0195 $CanB^{E19G}$ selected for growth in the absence of supplemental $CO_2$ with targeted insertion of $CanB^{19E}$ | This study |
| NY0195Δ*aasN* | NY0195 transformed with pDR46 to introduce a $Cm^R$ cassette and knockout *aasN* | This study |
| NY0195Δ*aasN* pDR48 | NY0195Δ*aasN* transformed with pDR48 with constitutive control of $AasN^{321Q}$ under the *opa* promoter | This study |
| NY0195Δ*aasN* pDR49 | NY0195Δ*aasN* transformed with pDR49 with constitutive control of $AasN^{321E}$ under the *opa* promoter | This study |
| FA1090Δ*canB* | FA1090 transformed with pDR35 to introduce a $Kan^R$ cassette and knockout *canB* | This study |

| | | |
|---|---|---|
| FA19 Cip Escapee 1 | FA19 selected for growth at 0.006 µg/mL ciprofloxacin, isolate 1 | This study |
| FA19 Cip Escapee 2 | FA19 selected for growth at 0.006 µg/mL ciprofloxacin, isolate 2 | This study |
| FA19 Cip Escapee 3 | FA19 selected for growth at 0.006 µg/mL ciprofloxacin, isolate 3 | This study |
| FA19 CanB$^{G19E}$ Cip Escapee 1 | FA19 CanB$^{G19E}$ selected for growth at 0.006 µg/mL ciprofloxacin, isolate 1 | This study |
| FA19 CanB$^{G19E}$ Cip Escapee 2 | FA19 CanB$^{G19E}$ selected for growth at 0.006 µg/mL ciprofloxacin, isolate 2 | This study |
| FA19 CanB$^{G19E}$ Cip Escapee 3 | FA19 CanB$^{G19E}$ selected for growth at 0.006 µg/mL ciprofloxacin, isolate 3 | This study |
| FA19 GyrA$^{91F}$ | FA19 selected for growth at 0.03 µg/mL $CO_2$ with targeted insertion of GyrA$^{91F}$ | This study |
| FA19 CanB$^{G19E}$ GyrA$^{91F}$ | FA19 CanB$^{G19E}$ selected for growth at 0.03 µg/mL $CO_2$ with targeted insertion of GyrA$^{91F}$ | This study |
| FA19 GyrA$^{91F/95G}$ | FA19 selected for growth at 0.03 µg/mL $CO_2$ with targeted insertion of GyrA$^{91F/95G}$ | This study |
| FA19 GyrA$^{91F/95G}$ | FA19 selected for growth at 0.03 µg/mL $CO_2$ with targeted insertion of GyrA$^{91F/95G}$ | This study |
| **Primers (underlined represents annealing sequence)** | | Template |
| DR_250_ngo2079_F | CCGCTGCTGCTGG | NG gDNA |
| DR_251_ngo2079_R | CCGCCGAACAGTCC | NG gDNA |
| DR_266_2079ko_1_F | TCCGTAGGTGAACCTGCGGGTGTCGCCGGTTTGTGT | NG gDNA |
| DR_272_2079ko_1_R | ATATTCTCATTTTAGCCATTTTGCCGTCCTCTGAAAAAGG | NG gDNA |
| DR_273_2079ko_2_F | CCTTTTTCAGAGGACGGCAAACGGGATCCGCCGTCT | AphA3 KanR cassette |
| DR_274_2079ko_2_R | CGCCGTACCGGTTTTTGTACGCGTCGACGCTTTTTA | AphA3 KanR cassette |
| DR_275_2079ko_3_F | TAAAAAGCGTCGACGCGTACAAAAACCGGTACGGCG | NG gDNA |
| DR_276_2079ko_3_R | CCGATTTTCTTCATATACCGTATATTTCAATCCGACTACG | NG gDNA |
| DR_277_2079ko_4_F | GATTGAAATATACGGTATATGAAGAAAATCGGACTGTTCG | NG gDNA |
| DR_271_2079ko_4_R | GCTAGTTATTGCTCAGCGGCACATTATGCATCGGGGCT | NG gDNA |

| | | |
|---|---|---|
| DR_194_Gibpuc_5_F | CCGCTGAGCAATAACTAGCGGATCCCCGGGTACCG | pUC19 |
| DR_195_Gibpuc_5_R | CCGCAGGTTCACCTACGGATCTAGAGTCGACCTGCAGG | pUC19 |
| DR_343_Cosel_1_F | TCCGTAGGTGAACCTGCGGGTGGCTTGCCGGCG | NG gDNA |
| DR_344_Cosel_1_R | CCTTGCATTCTAAAACCTTATAACGTTTCCGTCCGTTTTC | NG gDNA |
| DR_345_Cosel_2_F | GGACGGAAACGTTATAAGGTTTTAGAATGCAAGGAACAGT | NG gDNA |
| DR_346_Cosel_2_R | CGGTTCGCCCGGTGCTTCAGCGACGCGTCGACGCTTTTTA | NG gDNA |
| DR_347_Cosel_3_F | AGATGTCTAAAAAGCGTCGACGCGTCGCTGAAGCACCGGG | NG gDNA |
| DR_348_Cosel_3_R | AAAAAGGGTTCGCAATACCGTATATTTCAATCCGACTACG | NG gDNA |
| DR_349_Cosel_4_F | CGGATTGAAATATACGGTATTGCGAACCCTTTTTCAGAGG | NG gDNA |
| DR_350_Cosel_4_R | TAGTTATTGCTCAGCGGTCATGATGTTTCCATTCCTTCCG | NG gDNA |
| DR_359_530ko_1_F | TCCGTAGGTGAACCTGCGGCGAATCCCTGAAAAACACGC | NG gDNA |
| DR_360_530ko_1_R | CACACGCTGACCGCATTCAAACTTCCTTTATGGTTGCGGA | NG gDNA |
| DR_361_530ko_2_F | GTCCGCAACCATAAAGGAAGTTTGAATGCGGTCAGCGTGT | pKH37 |
| DR_362_530ko_2_R | GGACGGGGCTTCGGACGGCACGGCGTTTACGCCCCGCCCT | pKH37 |
| DR_363_530ko_3_F | GCGATGAGTGGCAGGGCGGGGCGTAAACGCCGTGCCGTCC | NG gDNA |
| DR_364_530ko_3_R | TAGTTATTGCTCAGCGGACGCAGGAGCAGGATATTTTAAA | NG gDNA |
| DR_62_pKH37_AphA3_1 _F | CAGGAGCTAAGGAAGCTAAACGGGATCCGCCGTCTGAAC | AphA3 KanR cassette |
| DR_63_pKH37_AphA3_1 _R | CACCAATAACTGCCTTAAAAAAACGCGTCGACGCTTTTTAGACATC | AphA3 KanR cassette |
| DR_64_pKH37_AphA3_2_F | GATGTCTAAAAAGCGTCGACGCGTTTTTTTTAAGGCAGTTATTGGTG | pKH37 |

| | | |
|---|---|---|
| DR_65_pKH37_AphA3_2_R | GCTCCAATTGGCCCTATAGTGAGATCTGCTGATCCTCTCA | pKH37 |
| DR_66_pKH37_AphA3_3_F | GTTCAGACGGCGGATCCCGTTTAGCTTCCTTAGCTCCTG | pKH37 |
| DR_67_pKH37_AphA3_3_R | TGAGAGGATCAGCAGATCTCACTATAGGGCCAATTGGAGC | pKH37 |
| DR_214_pMR32_AphA3_1_F | ACCCCAGAGTCCCGCGCGAAACGATCCTCATCCTG | pMR32 |
| DR_215_pMR32_AphA3_1_R | AAAGCGTCGACGCGTGTTAAGGGATTAGCTCCTGAAAATC | pMR32 |
| DR_216_pMR32_AphA3_2_F | TTTCAGGAGCTAATCCCTTAACACGCGTCGACGCTTTTTA | AphA3 KanR cassette |
| DR_217_pMR32_AphA3_2_R | GGAGGAAAAAATAAAGAGGGTTATACGGGATCCGCCGTCT | AphA3 KanR cassette |
| DR_218_pMR32_AphA3_3_F | GATCCCGTATAACCCTCTTTATTTTTTCCTCCTTATAAAA | pMR32 |
| DR_219_pMR32_AphA3_3_R | CAGGATGAGGATCGTTTCGCGCGGGACTCTGGGGT | pMR32 |
| DR_306_ngo2079_PacI_F | TCTTTAATTAAATGAGCGAGTTGAGCGAAAT | NG gDNA |
| DR_280_ngo2079_flag_XbaI_R | TAGTATTCTAGAGACTACAAAGACGATGACGACAAGTGATGTTTCCATTCCTTCCGATAA | NG gDNA |
| DR_341_ngo0530_PacI_F | TCTGTATTAATTAAATGAACCGGACTTATGCCAAT | NG gDNA |
| DR_342_ngo0530_XbaI_R | TCTGTATCTAGATCATTTGTTTCCTTCAAACTGCTC | NG gDNA |
| DR_402_can_cPCR_F | GATATCCGTGGCAAGGACG | EC gDNA |
| DR_403_can_cPCR_R | CGTTTAGCGTTCACGACATCC | EC gDNA |
| DR_404_can_allex_1_F | GAGAGCTTAGTACGTTAAACATGAGAGCTTAGTACGTGAAACATGAG | pRE107 |
| DR_405_can_allex_1_R | CGACTTCTGGGCGCTGTCTAGAAGAAGCTTGGGATCCG | pRE107 |
| DR_406_can_allex_2_F | CGTCTGGAAGAGCTGTTTGTTTGAGCTCTCCCGGGAATTC | pRE107 |
| DR_407_can_allex_2_R | CTCATGTTTCACGTACTAAGCTCTCATGTTTAACGTACTAAGCTCTC | pRE107 |
| DR_408_can_allex_3_F | CGGATCCCAAGCTTCTTCTAGACAGCGCCCAGAAGTCG | EC gDNA |
| DR_409_can_allex_3_R | GGTTACAGGTCGTTAACCTCCAACTTCACACTTACTCGTCCAG | EC gDNA |

| | | |
|---|---|---|
| DR_410_can_allex_4_F | CTGGACGAGTAAGTGTGAAGTTGGAGGTTAACGACCTGTAACC | EC gDNA |
| DR_411_can_allex_4_R | GAATTCCCGGGAGAGCTCAAACAAACAGCTCTTCCAGACG | EC gDNA |
| DR_380_gyrA_F | TCCGTAGGTGAACCTGCGGATGGAGGCACGGGCA | NG gDNA |
| DR_383_FA19gyrA_R | GCTAGTTATTGCTCAGCGGTTCGCGTTCGCCGTTTT | NG gDNA |

**Supplementary Table 1**. Plasmids, strains, and primers used in this study.

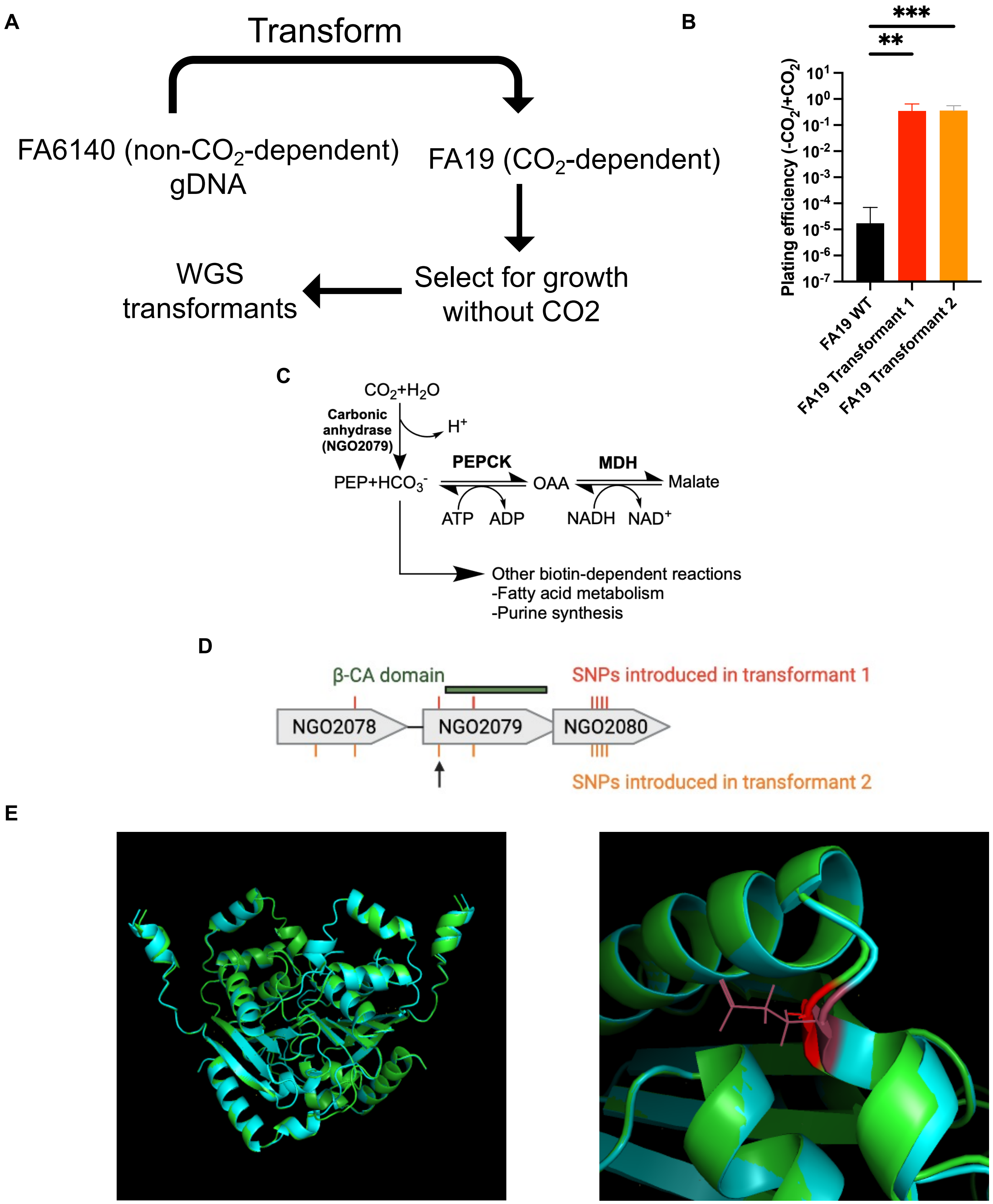
A
Transform
FA6140 (non-$CO_2$-dependent) gDNA
FA19 ($CO_2$-dependent)
Select for growth without CO2
WGS transformants
B
***
**
Plating efficiency (-$CO_2$/+$CO_2$)
$10^{1}$
$10^{0}$
$10^{-1}$
$10^{-2}$
$10^{-3}$
$10^{-4}$
$10^{-5}$
$10^{-6}$
$10^{-7}$
FA19 WT
FA19 Transformant 1
FA19 Transformant 2
C
$CO_2+H_2O$
Carbonic anhydrase (NGO2079)
$H^+$
PEP+$HCO_3^-$
PEPCK
OAA
MDH
Malate
ATP
ADP
NADH
$NAD^+$
Other biotin-dependent reactions
-Fatty acid metabolism
-Purine synthesis
D
β-CA domain
SNPs introduced in transformant 1
NGO2078
NGO2079
NGO2080
SNPs introduced in transformant 2
E

**Supplementary Figure 1**. Undirected genetic approaches identify a variant of CanB as causative for $CO_2$-dependence. **(A)** Schematic of undirected transformation of FA19 (parental CanB$^{19G}$) to identify causative factors for $CO_2$-dependence **(B)** Plating efficiency in the presence and absence of $CO_2$ following undirected transformation of gDNA from *N. gonorrhoeae* strain FA6140 into the *N. gonorrhoeae* lab strain FA19 (N ≥ 6 from two independent experiments). Significance determined by Mann-Whitney U test. **(C)** Putative reaction catalyzed by NGO2079/CanB and downstream metabolic products. **(D)** SNPs present in whole-genome sequencing of two transformants in **(B)**. Arrow indicates the SNP identified in **Figure 1**. **(E)** **(Left)** Alphafold predicted homodimeric structures of the CanB$^{19E}$ variant (teal) and the CanB$^{19G}$ variant (green). **(Right)** Magnified Alphafold predicted structures overlaying the glutamate (maroon) and glycine (red) at position 19.

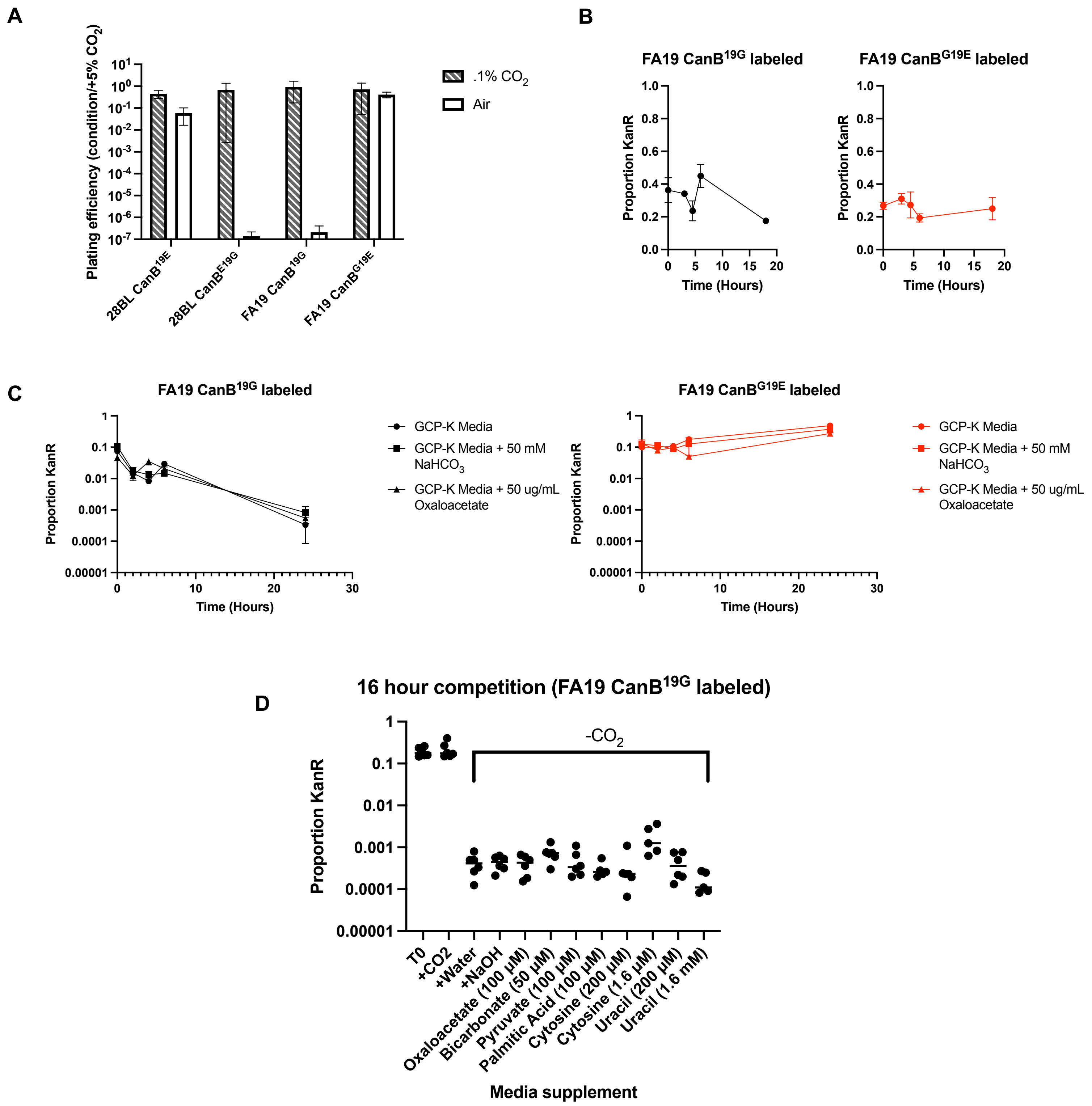

A
Plating efficiency (condition/+5% CO2)
.1% CO2
Air
28BL CanB19E
28BL CanBE19G
FA19 CanB19G
FA19 CanBG19E
B
FA19 CanB19G labeled
FA19 CanBG19E labeled
Proportion KanR
Time (Hours)
C
FA19 CanB19G labeled
FA19 CanBG19E labeled
GCP-K Media
GCP-K Media + 50 mM NaHCO3
GCP-K Media + 50 ug/mL Oxaloacetate
Proportion KanR
Time (Hours)
D
16 hour competition (FA19 CanB19G labeled)
-CO2
Proportion KanR
T0
+CO2
+Water
+NaOH
Oxaloacetate (100 μM)
Bicarbonate (50 μM)
Pyruvate (100 μM)
Palmitic Acid (100 μM)
Cytosine (200 μM)
Cytosine (1.6 μM)
Uracil (200 μM)
Uracil (1.6 mM)
Media supplement

**Supplementary Figure 2.** The CanB$^{19E}$ variant has an advantage in the absence of $CO_2$. **(A)** Plating efficiency at different $CO_2$ concentrations for *N. gonorrhoeae* strains 28BL (parental CanB$^{19E}$) and FA19 (parental CanB$^{19G}$) with isogenic CanB mutants (N ≥ 3, from two independent experiments). **(B)** Competition experiment in the presence of $CO_2$ between FA19 CanB$^{19G}$ and FA19 CanB$^{E19G}$ with (**left**) the FA19 CanB$^{19G}$ strain kanamycin-labeled and **(right)** the FA19 CanB$^{E19G}$ kanamycin-labeled. The proportion of colony forming units (CFUs) that are kanamycin resistant is graphed against time (N=3, representative of two independent experiments). **(C)** Similar to **(B)**, timecourse of competition between isogenic CanB FA19 strains in the presence of media supplementation and the absence of supplemental $CO_2$ (N=3, representative of two independent experiments). **(D)** As in **(C)**, competition between kanamycin-labeled FA19 CanB$^{G19E}$ and unlabeled FA19 CanB$^{19G}$ with selected metabolite supplementation assayed at 16 hours in the absence of supplemental $CO_2$ (N=6, from two independent experiments).

A

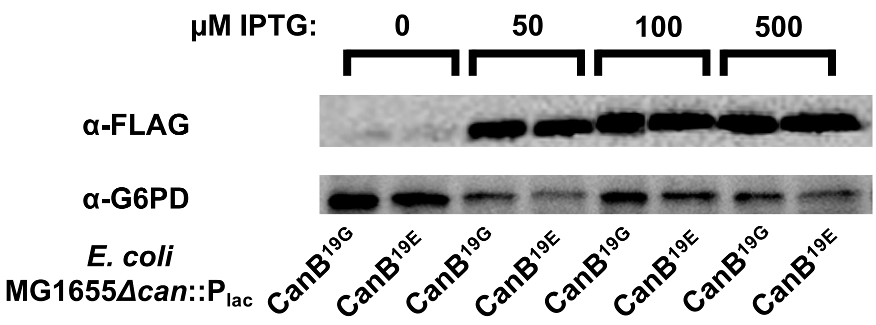
µM IPTG: 0 50 100 500
α-FLAG
α-G6PD
E. coli MG1655Δcan::Plac
CanB19G CanB19E CanB19G CanB19E CanB19G CanB19E CanB19G CanB19E

B

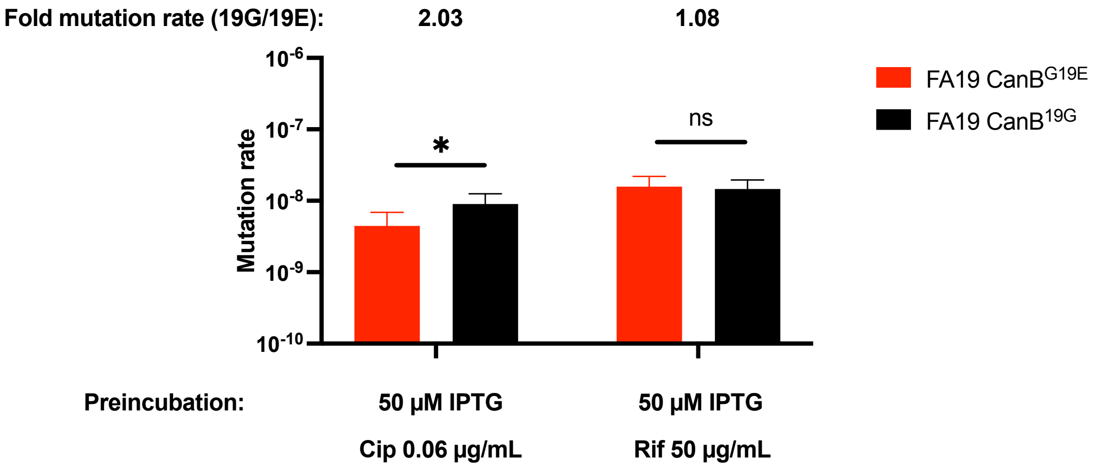
Fold mutation rate (19G/19E): 2.03 1.08
Mutation rate
10-6
10-7
10-8
10-9
10-10
*
ns
FA19 CanBG19E
FA19 CanB19G
Preincubation:
50 µM IPTG
Cip 0.06 µg/mL
50 µM IPTG
Rif 50 µg/mL

**Supplementary Figure 3.** In complemented *E. coli* MG1655*Δcan*, protein stability is similar between variants, and ciprofloxacin acquisition is associated with $CanB^{19G}$. **(A)** Western blot of *E. coli* MG1655*Δcan* complemented with IPTG-inducible CanB variants C-terminally FLAG-tagged with G6PD as loading control (representative of two independent experiments). **(B)** Fluctuation analysis of *E. coli* MG1655*Δcan* complemented with IPTG-inducible CanB variants and preincubated with 50 µM IPTG (N=180, from two independent experiments). Significance by unpaired two sample t-test. *$p<0.05$, **$p<0.01$, ***$p<0.001$

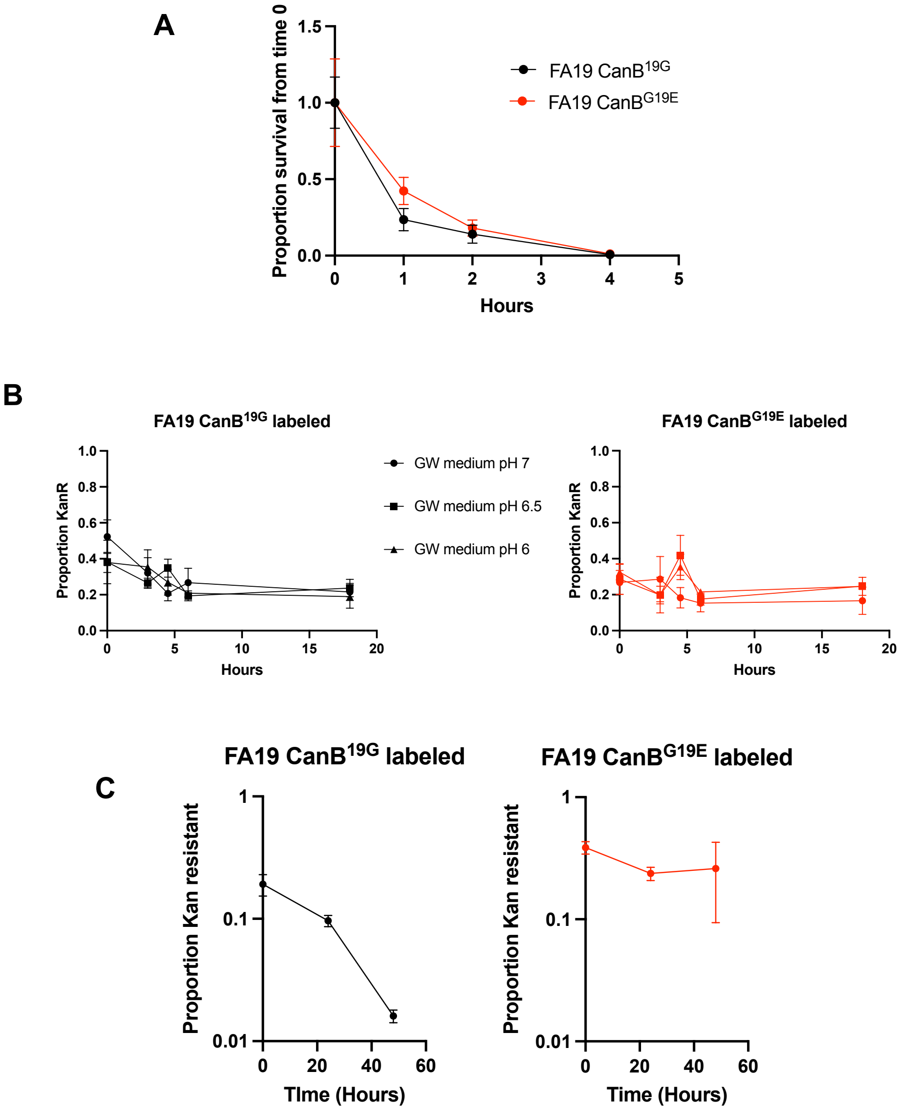
A
Proportion survival from time 0
1.5
1.0
0.5
0.0
0
1
2
3
4
5
Hours
FA19 CanB19G
FA19 CanBG19E
B
FA19 CanB19G labeled
Proportion KanR
1.0
0.8
0.6
0.4
0.2
0.0
0
5
10
15
20
Hours
GW medium pH 7
GW medium pH 6.5
GW medium pH 6
FA19 CanBG19E labeled
C
FA19 CanB19G labeled
FA19 CanBG19E labeled
Proportion Kan resistant
1
0.1
0.01
0
20
40
60
TIme (Hours)
Time (Hours)

**Supplementary Figure 4.** The 19E variant of NGO2079 is advantaged under anaerobic conditions, but not at moderately low pH or within macrophages. **(A)** Gentamicin intracellular protection assay in RAW 264.7 macrophages as measured by CFUs (N=3/timepoint, representative of two independent experiments). **(B)** Competition experiment as in **Supplementary Figure 2A** in pH-adjusted Graver-Wade media (N=3). **(C)** Competition under anaerobic conditions between FA19 isogenic CanB strains on GCB-K agar (N=3, representative of two independent experiments).

A

| Drug MIC | FA19 CanB$^{19G}$ | FA19 CanB$^{G19E}$ |
|---|---|---|
| Sulfamethoxazole | >256 μg/mL | >256 μg/mL |
| TMP/SMX | 2 μg/mL | 2 μg/mL |
| Trimethoprim | 4 μg/mL | 4 μg/mL |

| Drug MIC | 28BL CanB$^{19E}$ | 28BL CanB$^{E19G}$ |
|---|---|---|
| Sulfamethoxazole | >256 μg/mL | >256 μg/mL |
| TMP/SMX | 12 μg/mL | 6 μg/mL |
| Trimethoprim | 4 μg/mL | 4 μg/mL |

B

| Drug MIC | MG1655Δ*can*::$P_{lac}$-CanB$^{19G}$ + 50 μM IPTG | MG1655Δ*can*::$P_{lac}$ -CanB$^{19E}$ + 50 μM IPTG |
|---|---|---|
| Sulfamethoxazole | 256 μg/mL | 256 μg/mL |

C

| Drug | FA19 CanB$^{19G}$ | FA19 CanB$^{G19E}$ |
|---|---|---|
| Erythromycin | .19 μg/mL | .19 μg/mL |
| Chloramphenicol | 0.75 μg/mL | 0.75 μg/mL |
| Ciprofloxacin | .003 μg/mL | .003 μg/mL |
| Benzylpenicillin | 0.16 μg/mL | 0.16 μg/mL |
| Ceftriaxone | <0.002 μg/mL | <0.002 μg/mL |

D

| CipR mutant | Mutation | Ciprofloxacin MIC (μg/mL) |
|---|---|---|
| FA19 CanB$^{19G}$ Escapee 1 | *gyrA*$^{95N}$ | .032 |
| FA19 CanB$^{19G}$ Escapee 2 | *gyrA*$^{95N}$ | .023 |
| FA19 CanB$^{19G}$ Escapee 3 | *gyrA*$^{95G}$ | .032 |
| FA19 CanB$^{G19E}$ Escapee 1 | *gyrA*$^{95N}$ | .032 |
| FA19 CanB$^{G19E}$ Escapee 2 | *gyrA*$^{95N}$ | .047 |
| FA19 CanB$^{G19E}$ Escapee 3 | *gyrA*$^{95N}$ | .032 |
| FA19 CanB$^{19G}$ GyrA$^{91F}$ | *gyrA*$^{91F}$ | .047 |
| FA19 CanB$^{G19E}$ GyrA$^{91F}$ | *gyrA*$^{91F}$ | .047 |
| FA19 CanB$^{19G}$ GyrA$^{91F/95G}$ | *gyrA*$^{91F/95G}$ | .094 |
| FA19 CanB$^{G9E}$ GyrA$^{91F/95G}$ | *gyrA*$^{91F/95G}$ | .094 |

**Supplementary Table 2.** Antibiotic MICs for strains in this study. **(A)** MICs for sulfamethoxazole, trimethoprim, and trimethoprim/sulfamethoxazole for the FA19 and 28BL pairs isogenic CanB variants. **(B)** Sulfamethoxazole susceptibility of MG1655Δ*can* complemented with isogenic CanB variants and induced with IPTG. **(C)** MICs for clinically relevant antibiotics for FA19 CanB$^{19G}$ and FA19 CanB$^{E19G}$ as determined by E-test and agar dilution plating. **(D)** *gyrA* mutations for spontaneous ciprofloxacin escapees or clean mutants of FA19 CanB$^{19G}$ and FA19 CanB$^{19E}$, along with associated ciprofloxacin MIC.

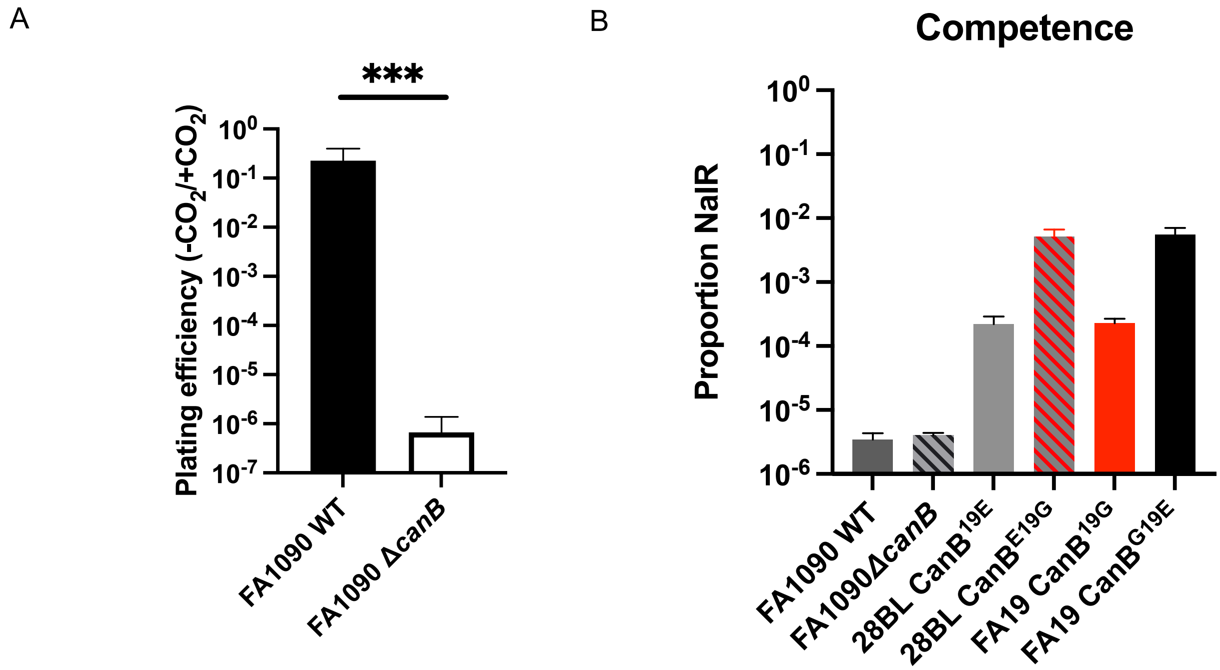
A
***
Plating efficiency (-CO2/+CO2)
10^0
10^-1
10^-2
10^-3
10^-4
10^-5
10^-6
10^-7
FA1090 WT
FA1090 ΔcanB
B
Competence
Proportion NalR
10^0
10^-1
10^-2
10^-3
10^-4
10^-5
10^-6
FA1090 WT
FA1090ΔcanB
28BL CanB19E
28BL CanBE19G
FA19 CanB19G
FA19 CanBG19E

**Supplemental Figure 5.** Loss of CanB does not appear to confer a loss of natural competence. **(A)** Plating efficiency in the presence and absence of $CO_2$ for a knockout of *CanB* in the *N. gonorrhoeae* lab strain FA1090 (N=6, from two independent experiments). Significance by Mann-Whitney U. **(B)** Natural competence of strains of *N. gonorrhoeae* as measured by uptake of a nalidixic acid resistance-conferring integrating plasmid (N=3, representative of two independent experiments). *p<0.05, **p<0.01, ***p<0.001

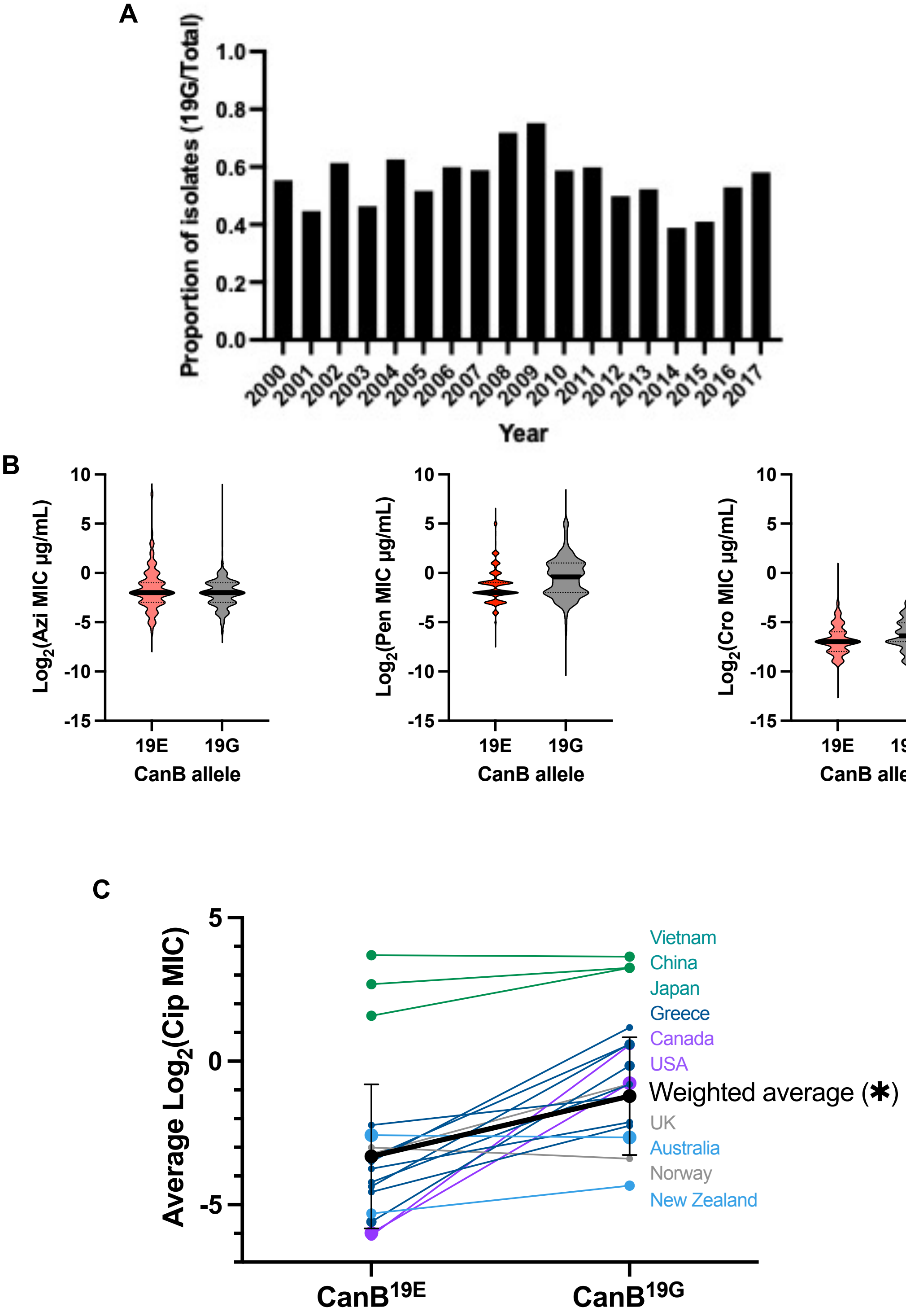
A
Proportion of isolates (19G/Total)
1.0
0.8
0.6
0.4
0.2
0.0
2000 2001 2002 2003 2004 2005 2006 2007 2008 2009 2010 2011 2012 2013 2014 2015 2016 2017
Year
B
Log2(Azi MIC μg/mL)
Log2(Pen MIC μg/mL)
Log2(Cro MIC μg/mL)
10
5
0
-5
-10
-15
19E
19G
CanB allele
C
Average Log2(Cip MIC)
5
0
-5
Vietnam
China
Japan
Greece
Canada
USA
Weighted average (✱)
UK
Australia
Norway
New Zealand
CanB19E
CanB19G

**Supplementary Figure 6.** The CanB$^{19G}$ variant is not strongly associated with higher resistance to multiple clinically relevant drugs, has remained relatively stable in proportion of sequenced isolates, and is associated with ciprofloxacin resistance across countries. **(A)** Proportion of sequenced isolates by year with the CanB$^{19G}$ variant. **(B)** Violin plots of drug MICs for ~10,000 *N. gonorrhoeae* clinical isolates. Black bar represents median, dotted lines represent $25^{th}$/$75^{th}$ percentiles. **(C)** Average $\log_2$ of ciprofloxacin MIC in µg/mL by country and by allele of CanB. Weighted average of all datasets is represented by black dots/line. Size of data point represents size of dataset. Error bars represent standard deviation as determined by weighted variance. Significance by unpaired two sample t-test. *$p<0.05$, **$p<0.01$, ***$p<0.001$

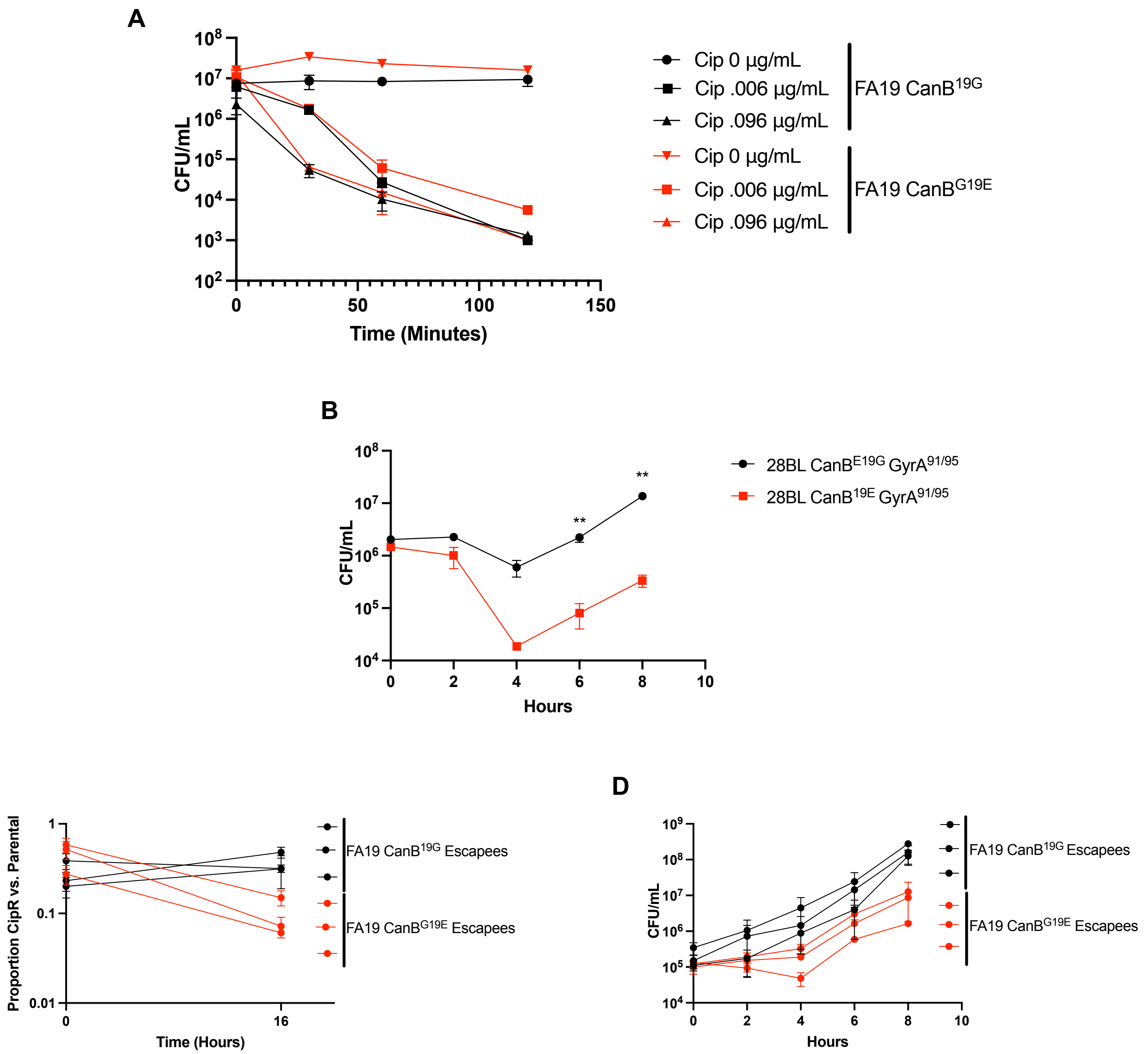
A
CFU/mL
Time (Minutes)
Cip 0 µg/mL
Cip .006 µg/mL
Cip .096 µg/mL
FA19 CanB19G
Cip 0 µg/mL
Cip .006 µg/mL
Cip .096 µg/mL
FA19 CanBG19E
B
CFU/mL
Hours
**
**
28BL CanBE19G GyrA91/95
28BL CanB19E GyrA91/95
C
Proportion CipR vs. Parental
Time (Hours)
FA19 CanB19G Escapees
FA19 CanBG19E Escapees
D
CFU/mL
Hours
FA19 CanB19G Escapees
FA19 CanBG19E Escapees

**Supplementary Figure 7.** The CanB$^{19G}$ variant does not affect killing by ciprofloxacin, but CanB$^{19G}$ provides an advantage in the presence of *gyrA* mutations across multiple *gyrA* alleles and strain backgrounds. **(A)** Kill curve of isogenic CanB FA19 at 2x MIC and 32x MIC (N=3, representative of two independent experiments). **(B)** Growth of CanB isogenic isogenic 28BL strains bearing ciprofloxacin resistance-determining GyrA$^{91F/95G}$ (N=3, representative of two independent experiments). Significance determined by unpaired two sample t-test. **(C)** Competition between spontaneously ciprofloxacin resistant isogenic CanB strains (see **Supplementary Figure 2**) and susceptible parental strains (N=3, representative of two independent experiments). **(D)** Growth curves of CanB isogenic FA19 strains with spontaneous ciprofloxacin resistance-determining *gyrA* alleles (N=3, representative of two independent experiments). *p<0.05, **p<0.01, ***p<0.001

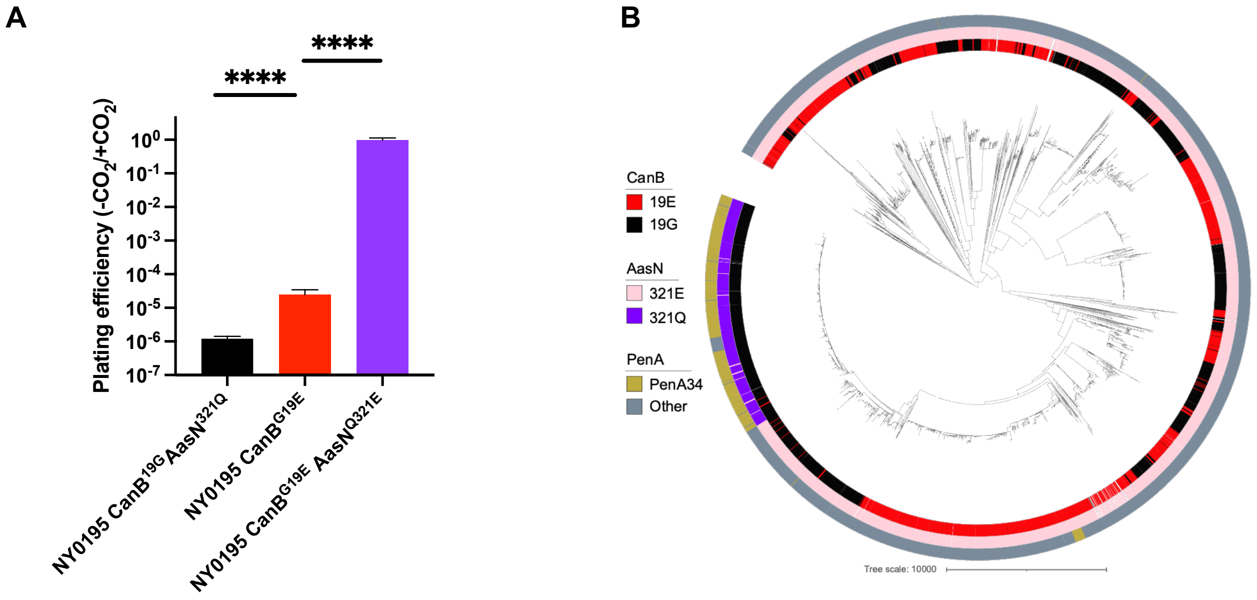

A
****
****
Plating efficiency (-CO2/+CO2)
10^0
10^-1
10^-2
10^-3
10^-4
10^-5
10^-6
10^-7
NY0195 CanB19G AasN321Q
NY0195 CanBG19E
NY0195 CanBG19E AasNQ321E
B
CanB
19E
19G
AasN
321E
321Q
PenA
PenA34
Other
Tree scale: 10000

**Supplementary Figure 8.** The relationship between CanB$^{19E}$ and $CO_2$-dependence depends on the acyl-ACP synthetase AasN in the PenA34 lineage of *N. gonorrhoeae*. **(A)** Plating efficiency in the absence and presence of supplemental $CO_2$ of clinical *N. gonorrhoeae* isolate NY0195 following the introduction of mutations in CanB and AasN (N=6, from two independent experiments). Significance by Mann-Whitney U test. **(B)** Maximum-likelihood tree as in **Figure 1A** overlaid with tracks indicating AasN allele and the ceftriaxone resistance determinant mosaic PenA34 allele. $^{*}p<0.05$, $^{**}p<0.01$, $^{***}p<0.001$, $^{****}p<0.0001$

### Methods and Materials

#### Bacterial strains, culture conditions, and MIC measurements

Strains are presented in Supplementary Table 1. *N. gonorrhoeae* was cultured on GCB agar (Difco) with Kellogg's supplements[1] (GCB-K) at 37°C. Strains were grown at concentrations of $CO_2$ as indicated. Antibiotic susceptibility testing was performed on GCB agar with Kellogg's supplements via Etest (BioMerieux (azithromycin, ciprofloxacin, benzylpenicillin, ceftriaxone, trimethoprim/sulfamethoxazole) or agar dilution (for trimethoprim and sulfamethoxazole). For plating efficiency experiments, we made serial dilutions, then plated in parallel on GCB agar with Kellogg's supplements. For anaerobic growth on solid medium, *N. gonorrhoeae* strains were first grown overnight on GCB agar with Kellogg's supplement and additional supplementation with 1.2 mM $NaNO_2$. For anaerobic growth curves, the overnight cultures were resuspended and grown in pre-reduced GCP medium (15 g/L proteose peptone 3 (ThermoFisher), 1 g/L soluble starch (ThermoFisher), 4 g/L $K_2HPO_4$, 1 g/L $KH_2PO_4$, 5 g/L NaCl)[2] with Kellogg's supplement and 1.2 mM $NaNO_2$. Anaerobic conditions were ensured using an anerobic hood and a gas mix consisting of 80% $N_2$, 10% $CO_2$, and 10% $H_2$. Optical density was measured at 600 nm. *E. coli* strains were cultured in LB.

#### Genomics pipeline and GWAS

Genomic analysis was conducted as in published work[3]. Reads were downloaded from sources also described[3]. These reads were inspected using FastQC (v0.11.7) (https://www.bioinformatics.babraham.ac.uk/projects/fastqc/) and removed for poor base quality or if GC content was divergent from expected values (~52-54%). Reads were then mapped to the NCCP11945 genome (RefSeq accession: NC_011035.1) using BWA-MEM (v0.7.17-r1188)[4]. We deduplicated using Picard (v2.8.0) (https://github.com/ broadinstitute/picard). Samples with <70% reads aligned or with less than 40% coverage were identified with BamQC in Qualimap (v2.2.1)[5] and discarded. Pilon (v1.16)[6] was used to call variants with a mindepth of 10 and a minmq of 20 and pseudogenomes were generated from Pilon VCFs by including all PASS sites and alternate alleles with AF>0.9, while all other sites were set to "N" and samples with >15% of sites called as missing were excluded. *De novo* assemblies were created using SPAdes (v3.12.0, with 8 threads, the --careful flag, and paired end reads where available)[7]. Contigs were then filtered for quality (coverage >10X, length >500 bp, total genome size ~2.0-2.3 Mbp). Assemblies were annotated using Prokka (v1.13)[8]. Core genes were clustered using Roary (v3.12, using flags -z -e -n -v -s -i 92)[9] and core intergenic regions were clustered using piggy (v1.2)[10]. Gubbins (v 2.3.4)[11] was then used to construct a recombination-corrected phylogeny with the aligned pseudogenomes and the resulting phylogeny was visualized in iTOL (v6)[12]. Metadata, including year, resistance determinants, and available susceptibility data have been collected and published[3]. For country level data, isolates with MICs that were annotated as greater than or less than that value were set to that value.

For the determination of GWAS phenotypes, thirty strains of *N. gonorrhoeae* were streaked on GCB agar with Kellogg's supplements in the presence and absence of supplemental 5% $CO_2$. These plates were then scored on a binary phenotype of growth/no growth in the absence of $CO_2$. For the computational aspect of the regression-based GWAS, we used a linear mixed model with a random effect to control for possible confounding by population structure. This linear mixed-model GWAS was run using Pyseer (v1.2.0 with default allele frequency filters)[13] based on 57,691 unitigs generated from the genome assemblies using unitig-counter (https://github.com/johnlees/unitig-counter). The Gubbins recombination-corrected phylogeny was used to parameterize the population structure random effects model in Pyseer. The resulting unitigs were mapped to the WHO F strain reference genome (Genbank accession: GCA_900087635.2) edited to contain only one locus of 23S rRNA using BWA-MEM (modified parameters -B 2 and -O 3). Significant unitigs were annotated through Pyseer's annotation pipeline. Unitigs were then further analyzed in Geneious Prime (v2021.0, http://www.geneious.com).

**Undirected transformation of *N. gonorrhoeae* and determination of $CO_2$-dependent plating efficiency**

For identification of the genetic modulator of $CO_2$ dependence (determined to be NGO2079), the $CO_2$-dependent laboratory strain FA19 was transformed with genomic DNA from the laboratory strain FA1090, which is $CO_2$-independent. Similarly, for identification of an additional mediator of $CO_2$-dependency (determined to be *aasN*), clinical isolate NY0195 was transformed with genomic DNA from the clinical isolate GCGS0457 using the following procedure. Liquid transformations were performed according to previously published protocols[2,14]. Piliated *N. gonorrhoeae* were grown on GCB-K agar overnight. The colonies were then resuspended in GCP medium with Kellogg's supplement, 10 mM $MgCl_2$, and approximately 400 ng genomic DNA. These suspensions were incubated at 37°C for 10 minutes in the presence of 5% $CO_2$ then rescued on non-selective GCB-K agar for 4 hours at 37°C with 5% $CO_2$. The recovered transformants were then re-plated on GCB-K agar and incubated overnight at 37°C without supplemental $CO_2$. The process was carried out in parallel without gDNA as a control for spontaneous mutation. Finally, colonies were subcultured for further analysis. To determine capacity for natural competence, strains of *N. gonorrhoeae* were transformed with pSY6, a plasmid conferring an allele of GyrB that confers nalidixic acid resistance[15]. Finally, serial dilutions of the transformants and negative control were plated in parallel on GCB-K agar plus Kellogg's supplement either without antibiotics or with 1 µg/mL nalidixic acid.

To determine the plating efficiency in the presence and absence of $CO_2$, *N. gonorrhoeae* was streaked overnight on GCB-K agar at 37°C with 5% $CO_2$. The resulting streaks were then serially diluted in at least triplicate and plated on GCB-K agar, with parallel dilutions being grown in the presence and absence of 5% $CO_2$ at 37°C. Plating efficiency was defined as the quotient of CFUs in the absence of $CO_2$ and CFUs in the presence of $CO_2$.

**Sequencing and structural prediction**

Following undirected transformation, genomic DNA from transformants was isolated using the PureLink Genomic DNA Mini kit (Life Technologies). Genomic DNA libraries were prepared and sequenced by the Microbial Genome Sequencing Center (https://www.migscenter.com/) on a NextSeq 2000. Resulting paired-end reads were then merged with BBMerge (v38.84)[16] and trimmed with BBDuk (v1.0). Reads were then mapped to the parental genomic DNA assembly and variants predicted in Geneious 2021.0 using default settings. Signal sequences were predicted using SignalP 6.0[17] and dimeric structures were predicted using AlphaFold (via Colab, https://colab.research.google.com/github/deepmind/alphafold/blob/main/notebooks/AlphaFold.ipynb)[18].

The resulting structures were then visualized in PyMOL (v2.5) (https://pymol.org).

**CanB variant composition within *Neisseria***

Variant composition within *Neisseria* and closely related species at NGO2079/NEIS2004 was aggregated from multiple sources, including a previously published set of 12,111 *N. gonorrhoeae* isolates[19], PubMLST (https://pubmlst.org/, using NEIS2004 as reference, to obtain variant composition from *N. meningitidis*, *N. polysaccharea*, *N. cinerea*, *N. subflava*, *N. mucosa*, *N. lactamica* strains) and blastp (https://blast.ncbi.nlm.nih.gov/, using NGO2079 from *N. gonorrhoeae* isolate FA1090 as query, to obtain variant composition from *N. sicca*, *B. denitrificans*, *N. elongata*, *E. corrodens*). Genomes from strains representing a diverse set of species, listed in Figure 1C, were assembled as above or downloaded from Genbank and annotated 23S rRNA sequences were extracted from each genome. Each genome contained only a single annotataed 23S rRNA or all annotated 23S rRNA sequences were identical.

Finally, a phylogenetic tree was constructed from these 23S rRNA sequences using RAxML (v8, 100 replicates, GTRGAMMA nucleotide model, Rapid Bootstrapping algorithm)[20]. The resulting tree was visualized in iTOL.

**Targeted transformation of CanB and AasN**

A ~1 kb PCR fragment surrounding the specific SNP of interest in *canB* was amplified using the primer pair (DR_250/DR_251) (see **Supplementary Table 1**). For *aasN*, the specific SNP of interest in *aasN* was amplified using primer pair (DR_341/DR_342). $CO_2$-dependent strains were transformed with approximately 600 ng of resulting PCR product and transformants were selected for growth on the absence of $CO_2$ as above. For selection of $CO_2$-dependent transformants from $CO_2$-independent parental strains, liquid transformations were performed as above and colonies were plated overnight on non-selective GCB-K agar. Colonies were then replica plated onto GCB-K agar and grown in parallel in the presence and absence of 5% $CO_2$. Finally, colonies that appeared to have $CO_2$-dependence were subcultured for further analysis. Transformation reactions using the alternative *canB* variant were used as a negative control to account for spontaneous mutation. For kanamycin co-selection, pDR50/51 was constructed using the following primer pairs: (DR_343/DR_344), (DR_345/DR_346), (DR_347/DR_348), (DR_349/DR_350), and (DR_194/DR_195) using Gibson assembly[21]. Liquid transformations were then performed on the above, followed by selection on GCB-K agar supplemented with 70 µg/mL kanamycin. Transformations without DNA were used as negative controls. All transformants were then checked by Sanger sequencing.

To knockout CanB, pDR35 was constructed using the following primer pairs: (DR_266/DR_272), (DR_273/DR_274), (DR_275/DR_276), (DR_277/DR_271), and (DR_194/195) using Gibson assembly and transformed as above. Similarly, to knockout AasN, pDR46 was constructed using primer pairs (DR_359/DR_360), (DR_361/DR362),

(DR_363/DR_364), and (DR_194/DR_195). pDR46 was then transformed into *N. gonorrhoeae* followed by selection on 8 µg/mL chloramphenicol in GCB-K agar and checked by Sanger sequencing. Finally, for complementation of AasN, pDR75, a kanamycin resistant derivative of pMR32 was cloned, using primer pairs (DR_214/DR_215), (DR_216/DR_217), and (DR_218/DR_219) and Gibson assembly. Complementation constructs pDR48 and pDR49 were then cloned from pDR75 using restriction enzyme cloning with PacI/XbaI and an insertion product amplified by primer pair (DR_341/DR_342). These products were transformed into the knockout strains, followed by selection on GCB-K agar supplemented with 70 µg/mL kanamycin.

**Competition of CanB variants**

Both FA19 $CanB^{19G}$ and $CanB^{G19E}$ were transformed with pDR1, a kanamycin cassette derivative (constructed with primer pairs (DR_62/DR_63), (DR_64/DR_65), and (DR_66/DR_67)) of pKH37[22]. The resulting transformants were selected on GCB-K agar supplemented with 50 µg/mL kanamycin and the resulting transformants were checked by colony PCR using primer pair (DR_62/DR_63). During competition experiments, FA19 $CanB^{19G}$ was grown at a 1:1 starting ratio with kanamycin-resistant FA19 $CanB^{G19E}$, while in parallel FA19 $CanB^{G19E}$ was grown at a 1:1 starting ratio with kanamycin resistant FA19 $CanB^{19G}$. At each timepoint, cultures were serially diluted and plated on both GCB-K agar and GCB-K agar supplemented with 50 µg/mL kanamycin. Finally, dilutions on both plates were quantified and the proportion of kanamycin resistant bacteria was quantified at each timepoint.

**pH or antibiotic kill curves and determination of intracellular pH**

Strains were plated overnight on GCB-K agar then scraped after 16-20 hours into liquid GCP medium with Kellogg's supplement. The cells were spun down in a microcentrifuge (10,000 rpm for one minute) then resuspended in GCP medium with Kellogg's supplement. Cells were then diluted to an optical density of 1.00 and incubated in either pH-adjusted GCP medium with

Kellogg's supplement (or, if noted, Graver-Wade medium[23]) or GCP medium with the defined concentration of antibiotics and Kellogg's supplement at 37°C in the absence of $CO_2$ in triplicate.

Intracellular pH was determined by using the BCECF-AM dye (ThermoFisher). Strains of interest were grown overnight on GCB-K agar with Kellogg's supplements. After 16-20 hours, streaks were resuspended in liquid GCP medium with Kellogg's supplement and diluted to a starting OD of 0.2. These strains were then grown in parallel in the presence and absence of 5% supplemental $CO_2$ for four hours in triplicate. Cells were then washed in PBS, resuspended in PBS with 1 µg/mL BCECF-AM dye, and incubated for thirty minutes. Finally, cells were measured for excitation/emission at both 440nm/535nm and 485nm/535nm in a plate reader. The fluorescence at both excitation/emission wavelengths from a control with dye but no cells was then subtracted from the measured 440nm/535nm and 485nm/535nm values. Finally, as the 440nm/535nm value is a proxy for cell density, the final (485nm/535nm)/(440nm/535nm) ratio was compared to a normal curve created using laboratory strain FA19 in the presence of Graver-Wade medium at multiple pHs and in the presence of 30 µg/mL of the decoupler CCCP (Sigma).

**Intracellular macrophage survival**

RAW264 macrophages were plated overnight in a 12-well plate at ~1 million cells/well in DMEM with 4 mM L-glutamine, 1.25 mM sodium pyruvate, and 10% heat-inactivated FBS (ThermoFisher). Cells were then infected with the strains of interest at an MOI of ~100. The plate was then centrifuged at 1300 rpm for 4 minutes to promote adherence and was incubated for 1 hour at 37°C with 5% $CO_2$. Cells were then washed three times in DMEM and finally replated in the supplemented DMEM with 100 µg/mL gentamicin. At 0, 1, 2, and 4 hours, three wells for each bacterial strain were resuspended in 300 mL of 0.01% Triton-X-100 in sterile PBS by vigorous pipetting. The Triton-X-100 solution was incubated for ten minutes then vortexed vigorously, followed by dilution plating on GCB-K agar to determine the total intracellular bacterial burden.

**Knockout of *can* and complementation of CanB variants in *E. coli***

pDR55, a derivative of the *sacB* allelic exchange vector pRE107[24], was cloned with primer pairs (DR_404/DR_405), (DR_406/DR_407), (DR_408/DR_409), and (DR_410/DR_411), to insert into *can* and transformed into the *E. coli* λpir DAP auxotroph β2155. The plasmid was then conjugated into *E. coli* MG1655 by incubating the two *E. coli* strains together at a 1:1 ratio on LB with 0.3 mM DAP in the presence of 5% supplemental $CO_2$. The resulting transformants were then plated onto LB without DAP in the presence of 5% supplemental $CO_2$. Finally, the resulting transformants were checked by colony PCR using primer pair (DR_402/DR_403) for successful knockouts. The resulting strains were then transformed with pDR42 and pDR43, both derived from pMR33[25] using restriction-enzyme cloning with enzymes PacI/XbaI and *canB*-FLAG amplified with primer pair (DR_280/DR_306), as well as empty-vector pMR33 and selected on LB agar with 20 µg/mL chloramphenicol. Finally, these strains were assessed for growth in the presence and absence of $CO_2$ at different levels of IPTG induction.

Proteins were quantified by Western blot. Briefly, MG1655Δ*can*::pDR42/43 was grown overnight in LB with 20 µg/mL chloramphenicol and 5% $CO_2$ at 37°C. Each strain was then diluted to OD 0.01 and then grown in the presence of different levels of IPTG inducer. After 4 hours, 1.5 mL of each culture was centrifuged (14,800 rpm for 3 minutes) and resuspended in lysis buffer (1% v/v Triton-X-100, 150 mM NaCl, 2 mM EDTA, 20 mM Tris-HCl pH 7.5, 1X v/v cOmplete protease inhibitor (Sigma). After 30 minutes of incubation at 4°C with agitation, lysed cells were centrifuged at 4°C (5,000 rpm for 5 minutes). The resulting supernatant was used for further protein analysis. Whole cell extracts were electrophoresed on 4-12 µg/mL Tris-Glycine gels

(Invitrogen) and transferred to a PVDF membrane with semi dry transfer. The membranes were then blocked in 5% milk, incubated with primary antibodies to G6PD (AssayPro) and FLAG-tag (ThermoFisher), then HRP-conjugated goat anti-rabbit secondary antibodies (ThermoFisher). Finally, the membranes were visualized with Pierce ECL Western Blotting Substrate (ThermoFisher) and visualized by chemiluminescent and colorimetric imaging.

**Fluctuation analysis and gyrase mutants**

*gyrA*$^{91F}$ and *gyrA*$^{91F/95G}$ alleles were amplified by PCR using primer pair (DR_380/DR_383) from clinical *N. gonorrhoeae* isolate NY0195. These alleles were then introduced into strains FA19 and 28BL using liquid transformation as described above and selected on GCB-K agar with .003 and .006 µg/mL ciprofloxacin respectively.

For fluctuation analysis, *N. gonorrhoeae* strains were first streaked overnight. The resulting streaks were then resuspended in liquid GCP with Kellogg's supplement, diluted to OD 0.1, then re-plated onto GCB-K agar with Kellogg's supplement either without antibiotics or with .001 µg/mL ciprofloxacin with 30 µl in each well of a 24-well plate (120 wells/condition). After 16-20 hours, each well was resuspended in 200 µl of liquid GCP and agitated with beads. From each well, 40 µl was plated onto overdried GCB-K agar with Kellogg's supplement and the antibiotic of interest. Colonies were then counted after 24 hours. Furthermore, six wells were used to determine the number of plated CFUs by serial dilution on nonselective GCB-K agar with Kellogg's supplement. Mutation rates were determined using the R package flan (v0.9, default conditions, plating efficiency 0.2)[26]. For *E. coli*, strains were grown in liquid overnight in LB supplemented with 20 µg/mL chloramphenicol in the presence of 5% $CO_2$. Each overnight was then re-diluted in 96-well plates to OD .0001 and grown in LB with 20 µg/mL chloramphenicol and 50 mM IPTG with 5% $CO_2$. After 16-20 hours, 40 µl from each well was plated onto overdried LB agar with the antibiotic of interest and 20 µg/mL chloramphenicol. Colonies were then counted after 24 hours. As with *N. gonorrhoeae*, six wells were used to determine the number of plated CFUs by serial dilution on LB agar with 20 μg/mL chloramphenicol. Mutation rates were calculated similarly to *N. gonorrhoeae*.
